# Matrix Stiffness Regulates Mechanotransduction and Vascular Network Formation of hiPSC-Derived Endothelial Progenitors Encapsulated in 3D Hydrogels

**DOI:** 10.1101/2025.04.11.648340

**Authors:** Jiwan Han, Kathleen Halwachs, Toni West, Bryce Larsen, Michael S. Sacks, Adrianne M. Rosales, Janet Zoldan

**Affiliations:** Department of Biomedical Engineering, The University of Texas at Austin, Austin, Texas; Department of Chemical Engineering, The University of Texas at Austin, Austin, Texas; James T. Willerson Center for Cardiovascular Modeling and Simulation, The Oden Institute for Computational Engineering and Sciences, Austin, TX, USA

**Keywords:** Hydrogel stiffness, Mechanotransduction, hiPSC-derived endothelial cells, Vascular morphogenesis, YAP/TAZ signaling

## Abstract

The mechanical properties of the extracellular matrix (ECM), particularly stiffness, regulate endothelial progenitor responses during vascular development, yet their behavior in physiologically compliant matrices (<1 kPa) remains underexplored. Using norbornene-modified hyaluronic acid (NorHA) hydrogels with tunable stiffness (190–884 Pa), we investigated how hydrogel stiffness influences cell morphology, endothelial maturation, mechanotransduction, and microvascular network formation in human induced pluripotent stem cell-derived endothelial progenitors (hiPSC-EPs). Our findings reveal a stiffness-dependent tradeoff between mechanotransduction and vascular network formation. At intermediate stiffness (551 Pa), cells exhibited the greatest increase in endothelial marker CD31 expression and Yes-associated protein (YAP)/ transcriptional coactivator with PDZ-binding motif (TAZ) nuclear translocation, indicating enhanced mechanotransduction and endothelial maturation. However, this did not translate to superior plexus formation. Instead, the most compliant matrix (190 Pa) supported greater vascular connectivity, characterized by longer branches (∼0.03/volume vs. 0.015 at 551 Pa) and enhanced actin remodeling. 3D cell contraction measurements revealed a 15.6-fold higher basal displacement in compliant hydrogels, suggesting that cell-generated forces and matrix deformability collectively drive vascular morphogenesis. Unlike prior studies focusing on pathological stiffness ranges (>10 kPa), our results emphasize that vascularization is not solely driven by the most mechanotransductive environment but rather by a balance of compliance, contractility, and cell-induced remodeling. These findings underscore the need to design hydrogels that provide sufficient mechanotransduction for endothelial maturation while maintaining compliance to support dynamic vascular morphogenesis. This work provides a mechanically tuned framework for optimizing microenvironments to balance endothelial differentiation and vascular network formation in tissue engineering and regenerative medicine.

## 1. Introduction

The mechanical properties of the extracellular matrix (ECM) regulate cellular behavior and fate determination during tissue development and disease progression [1-3]. In vascular biology, matrix stiffness modulates endothelial cell phenotype through alterations in cell morphology, mechanotransduction signaling, and the expression of endothelial markers including CD31 and CD34 [4-6]. During vascular development, endothelial cells encounter a range of matrix stiffnesses, transitioning from the capillaries (∼3.6 kPa) through surrounding tissues (∼0.27 kPa) to form microvascular networks in matrices of 33.59-202 Pa [7-9]. This mechanical environment influences microvascular development and the formation of new capillary networks. Understanding these mechanical parameters advances therapeutic angiogenesis strategies, as the establishment of functional microvasculature remains a significant challenge in treating ischemic diseases and engineering vascularized tissues [3, 10, 11].

Mechanistic studies using human umbilical vein endothelial cells (HUVECs) have established that endothelial cells respond to substrate mechanics through defined mechanotransduction pathways [12-14]. On substrates exceeding 10 kPa, HUVECs demonstrate enhanced spreading, increased stress fiber formation, and elevated inflammatory responses [15-17]. These mechanical responses characterize pathological conditions such as atherosclerosis, where arterial wall stiffening alters endothelial function through mechanotransduction signaling [18, 19]. Two-dimensional culture studies with primary endothelial cells have demonstrated that increased substrate stiffness modifies endothelial barrier function, inflammatory activation, and angiogenic capacity [20, 21]. However, 2D systems do not replicate the three-dimensional architecture of native vasculature, where cells experience omnidirectional mechanical cues that elicit distinct morphological and functional responses [22-24].

In three-dimensional (3D) systems, a wide range of environmental parameters have been investigated for their effects on endothelial cell behavior. These studies have employed diverse materials (fibrin, collagen, hyaluronic acid and various composites) with widely varying densities and properties [25-28]. Furthermore, researchers have utilized HUVECS, primary endothelial cells, and human induced pluripotent stem cell-derived endothelial progenitors (hiPSC-EPs) in these various 3D environments [25-28]. However, in most cases, these variables are not systematically decoupled; for example, the material choice and changing of matrix densities alters stiffness, confounding interpretation of mechanosensitive responses [26]. We have previously described a hydrogel platform that enables independent tuning of stiffness without altering composition, and similar approaches have since been used to study how stiffness alone influences hiPSC-EC phenotypes [29-31]. However, data remain limited on endothelial progenitor cell mechanotransduction and differentiation within physiologically compliant 3D matrices (<1 kPa) that mimic early microvascular environments.

Here, we selected hyaluronic acid for hydrogel formulation due its abundance in the extracellular matrix and its key roles in cellular processes including migration, proliferation, angiogenesis, and morphogenesis [32-36]. Functionalization of hyaluronic acid with synthetic crosslinking groups allows hydrogel formation with highly tunable mechanics that mimic those of native ECM. Norbornene-modified hyaluronic acid (NorHA) hydrogels in particular offer specific advantages for investigating mechanobiological questions in three-dimensional environments [37]. For example, reaction of the norbornenes via thiol-ene chemistry enables cell attachment via the incorporation of adhesive motifs and local remodeling of encapsulated cells via the use of proteolytically degradable peptide crosslinkers [38]. Furthermore, the high selectivity of the thiol-ene reaction allows precise control over hydrogel crosslinking without biological cross-reactivity, enabling systematic investigation of mechanical effects on cell behavior [38].

The Yes-associated protein (YAP) and transcriptional coactivator with PDZ-binding motif (TAZ) function as primary mechanotransduction mediators across multiple cell types, including both mature endothelial cells and endothelial progenitors [40, 41]. These factors regulate transcriptional responses through mechanically-induced nuclear-cytoplasmic shuttling [2]. In primary endothelial cells, YAP/TAZ nuclear localization increases with matrix stiffening (>15 kPa) during pathological processes including atherosclerosis and tumor angiogenesis [42, 43]. However, the mechanisms governing endothelial specification within physiologically soft matrices remain undefined [44, 45]. Specifically, the temporal correlation between YAP/TAZ activation and endothelial marker expression (CD31/CD34) in 3D environments requires systematic investigation with endothelial progenitor populations.

The relationship between matrix mechanics and endothelial cell phenotype has implications for vascular pathologies and therapeutic strategies. Matrix stiffening in fibrotic conditions and solid cancer tumors disrupts vascular network formation, leading to altered tissue perfusion [46-48]. In solid tumors, however, this stiffening enhances YAP/TAZ nuclear localization in response to mechanical cues, as demonstrated by Scott et al. [49], who showed that stiffness, dimensionality, and cell shape collectively drive these mechanotransduction outcomes. This activation can amplify angiogenic signaling, such as through the VEGF-YAP/TAZ axis known to promote pathological vascularization in tumors [45, 49]. These findings highlight the critical, context-dependent role of matrix mechanics in regulating endothelial function and vascular development. In tissue engineering applications, the mechanical properties of the matrix environment significantly influence vascular network formation [50]. Understanding the precise mechanical requirements for endothelial specification and network formation can inform the design of therapeutically relevant tissue engineering strategies.

Here, we investigate the effects of matrix stiffness on endothelial cell differentiation and mechanotransduction using human induced pluripotent stem cell-derived endothelial progenitors (hiPSC-EPs). Unlike previous studies that primarily focus on HUVECs and primary endothelial cells, hiPSC-EPs offer distinct advantages for studying developmental mechanobiology, as they represent an early stage of vascular development while maintaining differentiation plasticity [51]. Using NorHA hydrogels with defined mechanical stiffnesses, we examined cellular responses across a physiologically relevant stiffness range of 190–884 Pa, mimicking the mechanical environment of early microvascular development [20, 52, 53]. This system allows us to probe endothelial progenitor behavior under mechanically distinct yet biologically relevant conditions [26, 54-56]. By analyzing changes in actin architecture, mechanotransduction, and basal cell contractility across physiologically relevant stiffness levels, we elucidate how hydrogel stiffness influences endothelial progenitor maturation and vascular morphogenesis across different time points. Our findings demonstrate that while intermediate stiffness promotes mechanotransduction, compliant matrices facilitate sustained cytoskeletal remodeling and higher basal contractility, ultimately leading to greater microvascular plexus interconnectivity over time. These results provide a framework for optimizing the development of angiogenic biomaterials that balance endothelial differentiation with functional vascular morphogenesis, advancing strategies for tissue engineering and regenerative medicine.

## 2. Materials and Methods

### 2.1 Synthesis of Norbornene-Modified Hyaluronic Acid (NorHA)

Norbornene-modified hyaluronic acid (NorHA) was synthesized similarly to what was described by Crosby et al. [29]. Briefly, sodium hyaluronic acid (Na-HA, 72 kDa, Lifecore) was first converted to its tetrabutylammonium salt (HA-TBA) by dissolving Na-HA in deionized water at 2 wt% with Dowex 50W ion exchange resin (4:1 resin:HA ratio by weight) under constant stirring at 600 RPM for 5 hours [57]. After filtering out the resin, the solution was titrated to pH 7.05 with tetrabutylammonium hydroxide, then frozen at - 80°C, and lyophilized. The resulting HA-TBA was dissolved in anhydrous DMSO (at a concentration of 20 mg/mL) under argon at 45°C with 5-norbornene-2-carboxylic acid (5-NB-2-CA) (7.5:1 M ratio of 5-NB-2-CA to HA-TBA repeat unit) and 4-(dimethylamino)pyridine (DMAP) (3.75:1 M ratio of DMAP to HA-TBA repeat unit). Next, di-tert-butyl dicarbonate (Boc2O) (1:1 M ratio of Boc2O to HA-TBA repeat unit) was added to the flask via syringe and the reaction was allowed to proceed for 24 hours. The reaction was quenched with four times the reaction volume of cold deionized water (4°C) and dialyzed for 3 days (Spectra/Por, MWCO: 6-8 kDa). After being removed from dialysis, sodium chloride (1 g per 100 mL) was dissolved in the solution, and the solution was drop-wise precipitated in 6X cold acetone (−20°C). The precipitate was collected by centrifugation, redissolved in deionized water, dialyzed for an additional week, frozen at -80°C, and lyophilized.

The degree of norbornene functionalization (∼52.5%) was confirmed by ^1^H NMR spectroscopy (Supplementary Fig. 1).

### 2.2 Peptide Synthesis and Purification

Following a similar protocol reported by Crosby et al. [29], RGD (GCGYGRGDSPG) and enzymatically degradable (DGD) peptide (KCGPQGIWGQCK) were synthesized by a Prelude X automated peptide synthesizer (Gyros Protein Technologies) with Rink Amide polystyrene resin (0.48 mmol/g, Chem Impex) as the solid support. The synthesis followed the standard Fmoc-based solid-phase peptide protocols. Upon completion of the synthesis, the peptides were cleaved from the resin using 10 mL of cleavage solution. The RGD cleavage cocktail was comprised of 94:2.5:2.5:1 trifluoroacetic acid:ethane-1,2-dithiol:water:triisopropylsilane solution, and the DEG cleavage cocktail was comprised of 92.5:2.5:2.5:2.5 trifluoroacetic acid:ethane-1,2-dithiol:water:triisopropylsilane solution. After mixing the resin with the cleavage cocktail for four hours, the resin was filtered out, and the peptides were precipitated into cold ether (−20°C, 10 fold volume) thrice and pelleted via centrifugation (0°C, 3000 RCF, 15 minutes). The RGD peptide was re-dissolved at a concentration of 10 mg/mL in 10:90 acetonitrile:water with 0.1% trifluoroacetic acid and the DEG peptide was re-dissolved at a concentration of 10 mg/mL in 20:80 acetonitrile:water with 0.1 v% trifluoroacetic acid. The peptide solutions were filtered with a PTFE 0.45 µM filter. Then, the peptides were purified with a C18 column on a Dionex UltiMate 3000 UHPLC using a linear gradient of acetonitrile with 0.1 v% trifluoroacetic acid in water with 0.1 v% trifluoroacetic acid, for 25 min at a solvent rate of 10 mL/min. The purified peptides were frozen at -80°C, lyophilized, and their masses were verified via matrix-assisted laser desorption/ionization time-of-flight (MALDI-TOF) mass spectrometry (Bruker autoflex maX MALDI-TOF) (Supplementary Fig. 2).

### 2.3 Fabrication of NorHA Hydrogels with Tunable Stiffness for hiPSC-Derived Endothelial Progenitor Encapsulation

To prepare the NorHA hydrogel, lyophilized NorHA was dissolved in endothelial cell growth medium 2 (EGM-2, PromoCell) to achieve a final concentration of 10 mg/mL. This working solution was then mixed with encapsulation media containing lithium phenyl-2,4,6-trimethylbenzoylphosphinate (LAP, 0.025 wt%) as the photoinitiator, RGD peptide (2 mM), ROCK inhibitor (Y-27632, Selleck Chemicals), recombinant human vascular endothelial growth factor (VEGF165, Acro Biosystems), and an enzymatically degradable peptide (DGD, KCGPQGIWGQCK). To modulate hydrogel stiffness, the concentration of DGD peptide was adjusted to 25%, 50%, 75%, or 100% of the available norbornene groups functionalized on the NorHA backbone. For instance, for the 25% NorHA hydrogel, DGD was added to react with 25% of the available norbornene groups, resulting in a final peptide concentration of 1.534 mM. The mixture was thoroughly vortexed to ensure uniformity before further processing. Supplementary Table 1 describes the amounts and concentrations of each component in the hydrogel formulation.

To encapsulate cells, hiPSC-derived CD34 positive endothelial progenitors were suspended in the NorHA hydrogel mixture at a density of 0.8 million cells/mL. The cell-laden hydrogel mixture was then plated onto a glass-bottom well plate and exposed to UV light (365 nm, 10 mW/cm^2^) immediately for 50 seconds to initiate cross-linking [29]. This approach ensures that the cells are uniformly distributed within the 3D matrix during cross-linking.

### 2.4 Rheological Measurements

The mechanical properties of NorHA hydrogels were characterized using an HR-2 rheometer (TA Instruments) with an 8 mm parallel plate geometry and a UV-transparent quartz plate. Hydrogel precursor solution (20 μL) was pipetted onto the quartz plate, which was connected to a mercury lamp (Omnicure Series 1500) fitted with a 365 nm filter. The hydrogel precursor solution was crosslinked *in situ* via exposure to 365 nm light (10 mW/cm^2^, 50 s). To obtain the storage modulus (G’), time sweeps were performed at 1 rad/s and 1% strain. All tests were performed at room temperature with at least four independent samples per condition.

### 2.5 Swelling Ratio Measurements

To determine the degree of mass swelling (Q_m_) of the NorHA hydrogels, 50 μL NorHA hydrogels were fabricated by pipetting hydrogel precursor solution onto a silicone sheet and exposing the macromer solution to 365 nm light (10 mW/cm^2^, 50 s) from a mercury lamp (Omnicure Series 1500) connected to a collimator. Upon gelation, the hydrogels were transferred to a 24 well plate and 1.5 mL phosphate buffered saline (PBS) was added to each well. The hydrogels were equilibrated in PBS at 37 °C for three days. The swollen hydrogels were then weighed in a 1.5 mL Eppendorf microcentrifuge tubes to measure their wet mass (M_w_). Subsequently, the hydrogels were frozen at −80 °C overnight and lyophilized using a Labconco freeze-dryer. The lyophilized samples were then weighed again to measure their dry mass (M_D_). Q_m_ was calculated as the ratio between M_w_ and M_D_.

### 2.6 hiPSC Culture and Maintenance

Human induced pluripotent stem cells (hiPSCs, and tdTomato^+^ hiPSCs; WiCell iPS-DF-19-9-11T) were cultured in Essential 8 medium (E8; Gibco A1517001) on vitronectin-coated 6-well plates (VTN-N, Gibco A14700). Medium was replenished daily, and cells were passaged upon reaching 60-70% confluence.

### 2.7 Endothelial Differentiation

A modified endothelial differentiation protocol based on methods described by Jalilian and Raimes [58], with subsequent refinements implemented by our research group [29, 59]. hiPSCs were initially seeded onto Geltrex-coated 6-well plates (Gibco A1413201) at a density of 8,000 cells/cm^2^. The culture medium consisted of E8 supplemented with 10 µM ROCK inhibitor (Y-27632, Selleckchem). Differentiation was initiated on day 0 by transitioning to differentiation medium 1 (DM1), comprising DMEM/F12 (Dulbecco’s Modified Eagle Medium/Nutrient Mixture F-12) supplemented with insulin-deficient B-27 (1:50, Gibco A1895601), N2 (1:100, Gibco 17502048), Activin A (25 ng/mL; PeproTech), BMP4 (30 ng/mL; PeproTech), and BIO (150 nM; Selleckchem). Cells were maintained in DM1 through day 1. From days 2-4, cultures were maintained in differentiation medium 2 (DM2), consisting of DMEM/F12 supplemented with B-27, N2, SB 431542 (2 µM; Selleckchem), and VEGF (50 ng/mL; PeproTech). Culture medium was exchanged every 24 hours throughout the differentiation process. Supplementary Table 2 provides a summary of hiPSC-EP DM1 and DM2 formulation.

### 2.8 Isolation of CD34^+^ hiPSC-EPs by Flow Cytometry

To isolate CD34^+^ cells, differentiated hiPSC-EPs were collected on day 7 and enzymatically dissociated with Accutase (STEMCELL Technologies, 07920) at 37°C for 10 minutes. After centrifugation at 300g for 5 minutes, cells were resuspended in ice-cold sorting buffer consisting of DPBS (GE Healthcare, SH30028.03) supplemented with 2 mM EDTA and 0.5% BSA (Sigma-Aldrich, A8412-100ML). The cell suspension was stained with CD34-PE antibody (Miltenyi Biotec, 130-113-741) at 4°C for 10 minutes, then diluted with additional sorting buffer. To ensure a single-cell suspension, the labeled cells were passed through 35 µm cell strainers (Corning, 352235). CD34^+^ cells were then isolated using a Bio-Rad S3e cell sorter.

### 2.9 Collagenase Degradation Study

To quantify hydrogel degradation via collagenase, 30 µL hydrogels were fabricated on a silicone sheet and transferred to a 24 well plate. The hydrogels were swollen in 1.5 mL of PBS for three days at 37°C. The hydrogels were massed, and the PBS was removed and replaced with 1 mL of collagenase solution. Hydrogels were removed and massed at many timepoints to determine the remaining mass. The collagenase solution consisted of 500 units/mL collagenase type I (ThermoFisher Scientific) dissolved in buffer containing 50 mM tricine, 10 mM calcium chloride, and 400 mM sodium chloride. The collagenase was isolated from Clostridium histolyticum, and was reported to have an activity of 250 units per mg. The collagenase was removed and replaced with fresh solution every 24 hours.

### 2.10 Cell shape Analysis

To investigate the relationship between matrix stiffness and cell morphology, a detailed three-dimensional analysis of encapsulated hiPSC-EPs was performed. Cell surface area and volume were quantified using an in-house custom Python software by analyzing cell surface membrane expression visualized through Wheat Germ Agglutinin (WGA) staining. First, image stacks were preprocessed into 8-bit images and single cells were identified by the presence of a single Hoechst stained nucleus using ImageJ-Fiji (Fuji). The custom software then produced a mask of the cell by identifying the cell membrane and nuclear fluorescence boundaries in the 3D images by implementing binary thresholding, salt and pepper removal, small hole filling, binary closing; adding of the red and blue signal; and choosing the connected component that was nearest the center of the xy plane of the image through implementation of classes found in the scikit image library [60]. The software then produced a finite element mesh from the mask with Lewiner marching cubes, taubin smoothing, removal of T-vertices, hole closing, and setting the mean element side length to 0.5 μm from the meshlab package [61]. From these reconstructions, the software calculated total surface area by summing the areas of all mesh elements, and volume by integrating the space enclosed by the surface mesh. To ensure accurate measurements, cells in contact with neighboring cells or image boundaries were excluded from analysis. Measurements were performed on at least 12 isolated cells per condition across three independent experiments. This analysis enabled quantitative comparison of cell morphology across various matrix stiffness conditions.

### 2.11 Immunostaining and Fluorescent Imaging

Cell-laden hydrogels were fixed in 4% paraformaldehyde (PFA) for 15 minutes at room temperature, followed by two 10-minute washes with Dulbecco’s phosphate-buffered saline (DPBS) containing 300 mM glycine. Next, cells were permeabilized with 0.1% Triton X-100 in DPBS for 45 minutes at room temperature, followed by the same washing procedure. Non-specific binding was minimized by incubating samples in blocking buffer (1% bovine serum albumin and 0.1% Triton X-100 in DPBS) for 45 minutes at room temperature. Primary antibodies were diluted in blocking buffer and applied overnight at 4°C: anti-CD34 (Invitrogen PA5-85917, 1:200), anti-CD31 (Invitrogen MA5-13188, 1:200), anti-YAP1 (Proteintech 66900-1-Ig, 1:100), and anti-TAZ (Invitrogen 703032, clone 7H33L24, 1:100). After four 10-minute washes with blocking buffer, samples were incubated overnight at 4°C with fluorophore-conjugated secondary antibodies diluted in blocking buffer: Alexa Fluor 488 (Invitrogen A-11001, 1:500) for CD31 and TAZ visualization, and Alexa Fluor 633 (Invitrogen A-21070, 1:500) for CD34 and YAP1 detection. The solution also contained DAPI (1:10,000) for nuclear visualization and rhodamine phalloidin (Invitrogen R415, 1:40) for F-actin labeling. Images were acquired using a Nikon W1 spinning disk confocal microscope equipped with a 60× water-immersion objective (Plan Apo VC 60×A WI DIC N2, NA 1.2). Z-stack images were acquired at 1 μm intervals using confocal microscopy to capture the complete spatial distribution of all fluorescent markers throughout the cell volume.

### 2.12 F-Actin Quantitative Immunofluorescence

F-actin z-stack images were acquired at 1 µm intervals and processed using ImageJ. Using background subtraction and Otsu thresholding procedures, binary images were generated, which were then skeletonized for quantitative analysis. The built-in Skeleton Analysis plugin in ImageJ was used to quantify key structural parameters, including the number of branches, number of junctions, average branch length, and longest shortest path. Total cell volume was determined by summing the areas of segmented F-actin masks across z-stack slices, acquired at a fixed interval of 1 μm. F-actin fluorescence was processed in ImageJ-Fiji, where the outermost cell boundaries were delineated, and internal voids were filled using the ‘Fill Holes’ algorithm to compute the cross-sectional area per slice. Given the 1 μm slice thickness, the cumulative sum of these areas directly corresponds to the total cell volume (μm). The extracted stress fiber parameters were then normalized to cell volume, yielding number of branches per volume, number of junctions per volume, average branch length per volume, and longest shortest path per volume. This normalization allowed for accurate assessment of cytoskeletal remodeling independent of cell size differences.

### 2.13 Quantitative Immunofluorescence Analysis of Cell Maturation

To examine the relative expression of endothelial markers CD31 and CD34, F-actin expression was used to define cell boundaries. These boundaries were used to generate cell masks in ImageJ, which were then applied to isolate CD31 and CD34 signals specifically associated with cell membranes. Following background subtraction with a 50-pixel rolling ball radius to eliminate non-specific fluorescence, the total fluorescent signal area across all z-stack slices was measured. The CD31/CD34 area ratio was calculated to evaluate endothelial cell maturation status at both day 4 and day 7 timepoints, where higher ratios indicate more mature endothelial characteristics.

### 2.14 Quantitative Immunofluorescence Analysis of Mechanotransduction

The intracellular distribution of YAP and TAZ between the cytosol and the nucleus was also examined. Nuclear regions were defined using DAPI fluorescence to generate nuclear masks, while cellular boundaries were determined using F-actin staining. To isolate cytoplasmic regions, the nuclear masks were subtracted from the cell boundary masks in ImageJ. After applying the same background subtraction parameters (50-pixel rolling ball radius), the total area of YAP and TAZ fluorescent signals within both nuclear and cytoplasmic compartments across all z-stack slices was quantified. The relative distribution was expressed as the ratio of nuclear to cytoplasmic area for each protein at both day 4 and day 7 timepoints. For detailed protein localization analysis, the Pearson correlation coefficients between YAP/TAZ signals and either nuclear (DAPI) or cytoplasmic (F-actin) markers across multiple z-stack slices was also calculated to validate the compartmental distribution patterns observed in our ratio measurements.

### 2.15 3D Cell Contractile Displacements Methods and Analysis

3-dimensional cell displacements experiments and analysis were performed, based on related traction force microscopy studies [62] with the following improvements. NorHA gels were prepared as stated above while implementing tdTomato-expressing hiPSC-EPs, with the addition of 0.05% TDS 0.51 μm diameter dragon green beads (FSDG003 Bangs Laboratories) to the gel precursor mix. Immediately prior to imaging, the hydrogels were stained for 30 minutes in Hoechst following manufacturer’s directions, and then 2.9 mL phenol-free media was added to each sample dish for imaging. The samples were imaged on a Nikon AXR confocal equipped with a Tokai hit stage incubator set to 37 □C and a 40x silicone oil objective that was run by the NIS-elements software (Nikon) at the Center for Biomedical Research Support Microscopy and Flow Cytometry Facility at UT Austin (RRID: SCR_021756). Simultaneous 405 nm, 488nm, and 561 nm laser exposure was implemented to illuminate the nuclei, dragon-green fiducial marker beads, and the tdTomato-expressing hiPSC-EPs, respectively. A 147×147×122 μm field of view (FOV) was imaged with isotropic voxels of side length 0.29 μm. Imaged cells had a single nucleus and were alone in the FOV so that as many fiducial markers as possible could be imaged in the FOV. Once cells were imaged in their basal states, 100 μL of 120 μM cytochalasin-D (CytoD) in phenol-free media was added to the sample dish, making the final concentration of the dish 4 μM. Samples were incubated for 40 min at 37 □C, and then each cell was reimaged in its fully-relaxed state. 3D images of each color channel were first converted into individual 8-bit RGB tiff slice images with ImageJ-Fiji (Fuji). The red fluorescence images from the CytoD state were introduced into the in-house Python script described above was executed to produce highly accurate mathematical representations of the cell images. Concurrently, the green fluorescence images were processed using a software we have previously published [63], FM-Track, which was implemented to track the movement of the fiducial marker beads within the gel going from the fully-relaxed CytoD-treated state to the basal state so that basal contractility of the cells could be analyzed. The displacement information produced from the bead tracking was corrected for displacements caused by the movement of the microscope stage during image acquisition with multivariate adaptive regression splines and filtered to remove spurious beads. Resultant bead locations and displacements were employed to produce a Gaussian process regression model of cell/gel displacements that was then implemented to interpolate the movement of the CytoD cell surface mesh because of basal contractility. Unit vectors pointing out from the center of the cell were used to normalize the direction of contraction events at the surface of the cell, making negative values represent contraction and positive events represent isovolumetric-related relaxation. Graphed values of displacement were then multiplied by -1 to make contractile responses positive. Visualizations of the bead, gel, and cell movement were performed in ParaView 5.12.0 (Kitware). Graphs and related statistical tests were produced in Prism 10.3.0 (GraphPad).

### 2.16 Vessel-Like Network Analysis: Length and Connectivity

Network characteristics such as total length, connectivity, and vessel diameter were assessed using a previously established computational pipeline [29, 59]. In this approach, confocal z-stack images were processed by filtering and binarization in ImageJ. The resulting images were then analyzed using a MATLAB script that constructs a nodal graph, where capillary-like structures are represented as nodes (branch/endpoints) and links (vessels).

### 2.17 Statistical Analysis

Statistical analysis was conducted using a one-way variance test and post hoc Tukey’s multiple comparison tests using Prism 10 software (GraphPad). Relationships were deemed significant at a threshold of p<0.05. The data are presented as mean values ± standard error. Statistical tests were performed after Robust regression and Outlier removal (ROUT) at Q=1% (GraphPad).

## 3. Results

### 3.1 NorHA Hydrogels represent a unique ECM-mimicking system that allows modulation of stiffness without altering the hydrogel’s degradation profile

Before cell encapsulation, the acellular hydrogels, composed of norbornene-modified hyaluronic acid (NorHA) and crosslinked with a degradable peptide (KCGPQGIWGQCK), hereafter referred to as DGD, was characterized. The concentration of DGD was varied and the impact on the stiffness of the NorHA hydrogel was measured. Rheological analysis revealed that the hydrogel’s storage modulus increased proportionally with DGD crosslinked peptide concentration from 190.0 ± 33.32 Pa at 25% crosslinking to 884.0 ± 94.76 Pa at 100% crosslinking (Fig. 1A). Next, the swelling ratio of these hydrogels was characterized. The equilibrium swelling ratio of the tested hydrogels was inversely related to the extent of hydrogel crosslinking, decreasing linearly (*R*^2^ = 0.99) from a wet/dry ratio of 54 at 25% crosslinking concentration to 35 at 100% crosslinking. (Fig. 1B). Moreover, when, the rate of NorHA hydrogel degradation by collagenase was tested for dependence on the integrity of the crosslinked peptide, which harbors a collagenase substrate (GPQGIWGQ), all hydrogel formulations maintained approximately 100% of their initial mass throughout the 96-hour observation period (±20%; Fig. 1C). This stability, observed regardless of crosslinking density, demonstrates that the primary NorHA network structure may be established through a combination of physical NorHA crosslinking (e.g., chain entanglement) and thiol-ene photoclick crosslinking of norbornene groups.

**Figure 1:**
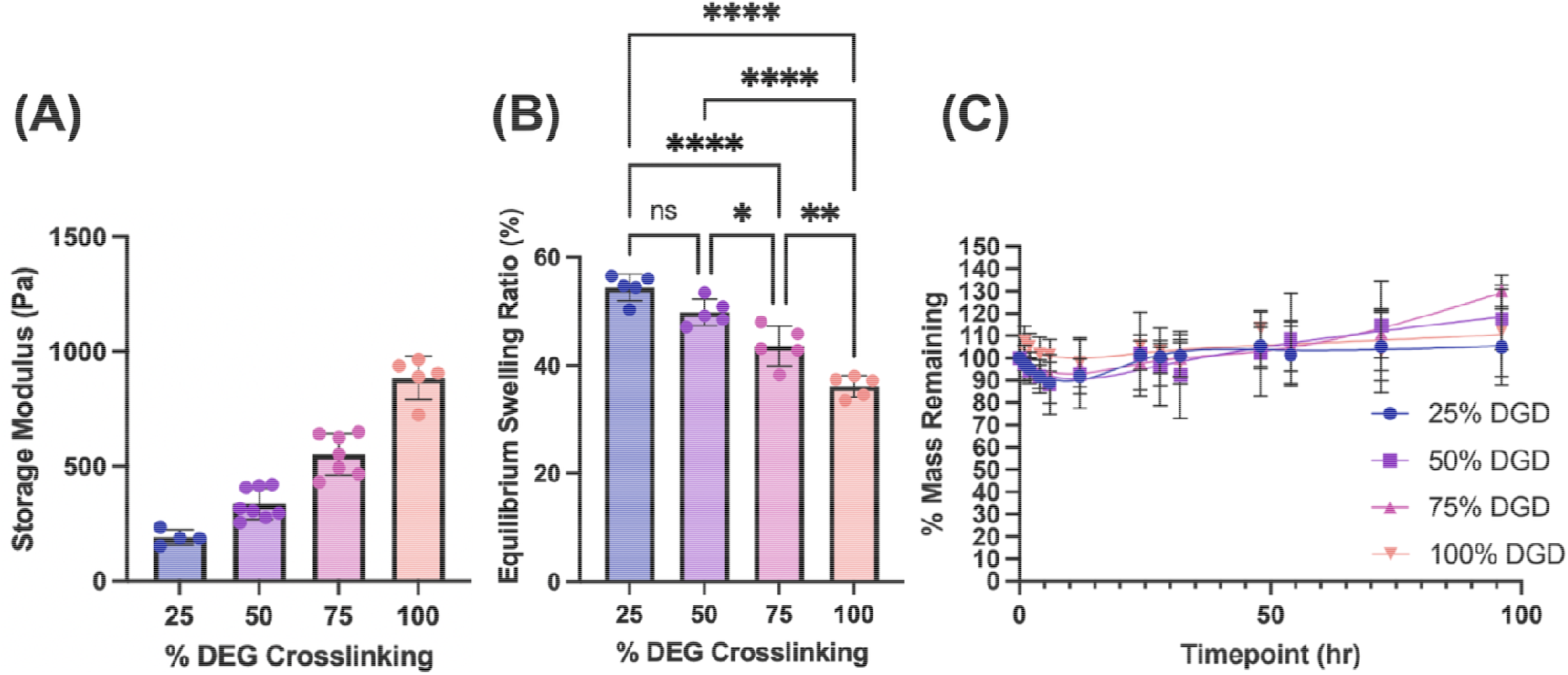
Charaterization of norbornene-modified hyaluronic acid (NorHA) hydrogels crosslinked with varying concentrations of degradable peptide (DGD, sequence: KCGPQGIWGQCK). **(A)** Storage modulus measurements showing increased hydrogel stiffness with higher DGD crosslinking percentages, **(B)** Equilibrium swelling ratio demonstrating an inverse relationship with crosslinking density **(C)** Mass retention profiles during collagenase degradation over 96 hours. Statistical significance is indicated as: *P ≤ 0.05, **P ≤ 0.01, ****P ≤ 0.0001.

Collectively, these measurements show that the stiffness of the NorHA hydrogel can be modified reproducibly and predictably within a physiologically relevant range [7-9] through varying the crosslinking density, while maintaining the structural integrity of the primary NorHA network and without changing the hydrogel’s chemical components. This system enabled a systematic investigation of how hydrogel stiffness influences cell morphology, differentiation markers, and mechanotransduction events.

### 3.2 The shape of hiPSC-EPs is nonlinearly dependent on NorHA stiffness

Once acellular hydrogel characterization was finalized, hiPSC-EPs were encapsulated in these hydrogel formulations and the cell-laden hydrogels were cultured for 4 days to examine how cell morphology varies with matrix stiffness. This was achieved through fluorescent labeling of the cell cytosol with F-actin and the nucleus with DAPI, followed by imaging and 3D cell surface reconstruction (Fig. 2). Representative 3D reconstructions of cell morphology and further quantification of cell volume and surface area reveal a pronounced dependence on hydrogel stiffness. hiPSC-EPs encapsulated in hydrogels with 190-551 Pa storage modulus were spindle-shaped. Notably, cells encapsulated in the 551 Pa hydrogels exhibited multiple surface protrusions. In contrast, cells encapsulated in the 884 Pa hydrogels were the most compact among all stiffness levels, appearing round with few cellular protrusions. Further quantifying cell surface and volume showed that both these parameters exhibited a nonlinear trend with increasing hydrogel stiffness. Cells encapsulated in the 551 Pa hydrogels displayed the highest volume and surface area compared to those encapsulated in either lower or higher stiffness conditions (Fig. 2C, 2D). Cells encapsulated in the 884 Pa hydrogels exhibited the lowest volume and surface area, consistent with previous observations indicating restricted cell spreading at higher matrix stiffness [14, 22, 29].

**Figure 2:**
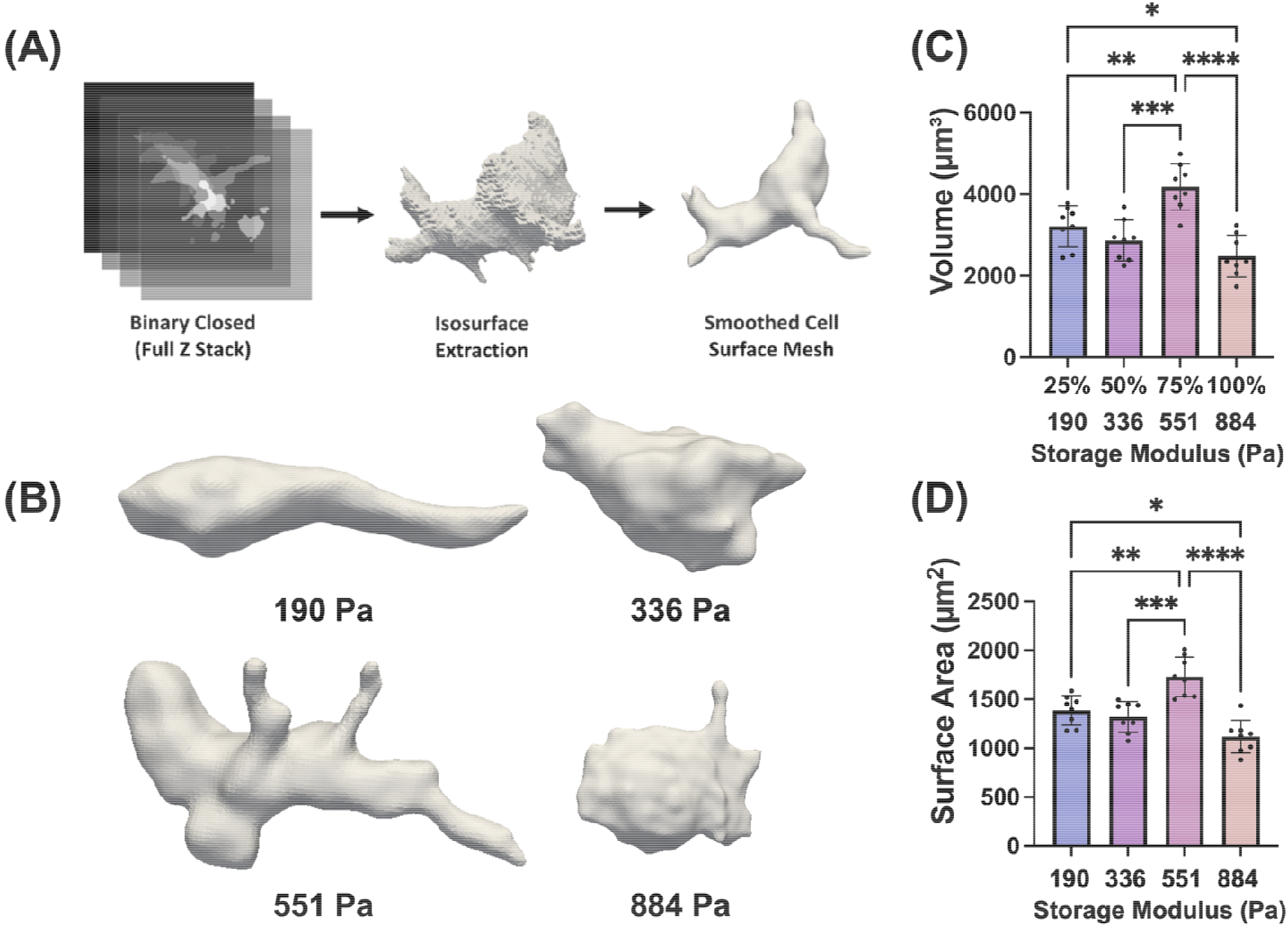
Three-dimensional morphological analysis of hiPSC-ECs in NorHA hydrogels of varying stiffness. **(A)** Workflow of cell surface reconstruction from Z-stack images. **(B)** Representative 3D reconstructions showing cell morphology at different storage moduli (190-884 Pa). **(C)** Cell volume and **(D)** surface area measurements. Statistical significance is indicated as: *P ≤ 0.05, **P ≤ 0.01, ***P ≤ 0.001, ****P ≤ 0.0001.

The impact of matrix stiffness on EC behavior has been reviewed by us and others [64, 65]. Overall, on 2D surfaces, ECs exhibit increased spreading and larger cell size on stiffer substrates, although most studies have focused on stiffness ranges more relevant to pathophysiological conditions. However, existing literature on the effect of stiffness on EC morphology and microvasculature formation in 3D microenvironments remains conflicting, largely due to the wide range of stiffness values studied, spanning from a few hundred Pascals [14, 23] to several thousand Pascals [66, 67]. Additionally, in 3D environments, stiffness is often confounded by other matrix properties such as degradability and diffusivity, further complicating the isolation of stiffness-dependent effects.

### 3.3 Actin Cytoskeleton Organization Depends on Matrix Stiffness

To better understand the morphological changes described above, the 3D F-actin network was reconstructed from its fluorescence channel across the Z-stack images of hiPSC-EPs encapsulated in hydrogels ranging from stiffness values of 190 to 884 Pa (Fig. 3). Quantifying the F-actin network revealed that while the number of branches per volume remained relatively consistent from 190 Pa to 551 Pa (varying by less than 10%), the junctions per volume increased by approximately 2.1-fold from 190 Pa (0.0025) to 551 Pa (0.0052), indicating enhanced cytoskeletal connectivity at intermediate stiffness levels. At 884 Pa, both parameters significantly decreased, with branches declining by 40% and junctions decreasing by 45% compared to 551 Pa, suggesting reduced actin polymerization and impaired cytoskeletal integrity. In contrast, average branch length per volume exhibited a stiffness-dependent decline, decreasing by ∼37% from 190 Pa (∼0.008) to 336 Pa (∼0.005) and by an additional 10% at 551 Pa (0.0045). Similarly, the longest shortest path per volume decreased by 33% from 190 Pa (0.03) to 336 Pa (0.02) and further declined by 25% at 551 Pa (0.015). These reductions in fiber length coincided with the significant increase in junctions per volume, suggesting that actin fibers became more interconnected rather than simply shorter. This trend indicates that as stiffness increased within this range, a denser and more branched cytoskeleton network was formed, leading to reduced individual fiber length due to increased branching points. At 884 Pa, however, both average branch length and longest shortest path per volume further declined by ∼33% and ∼20% respectively compared to 551 Pa, accompanied by a reduction in junctions, suggesting a shift from an interconnected cytoskeleton network to a more fragmented cytoskeletal structure. Collectively, these findings demonstrate that matrix stiffness modulates cytoskeleton organization in a biphasic manner, where intermediate stiffness (551 Pa) enhances cytoskeletal connectivity through increased actin junction formation, while excessive stiffness (884 Pa) leads to a loss of network integrity, reducing fiber elongation and connectivity.

**Figure 3:**
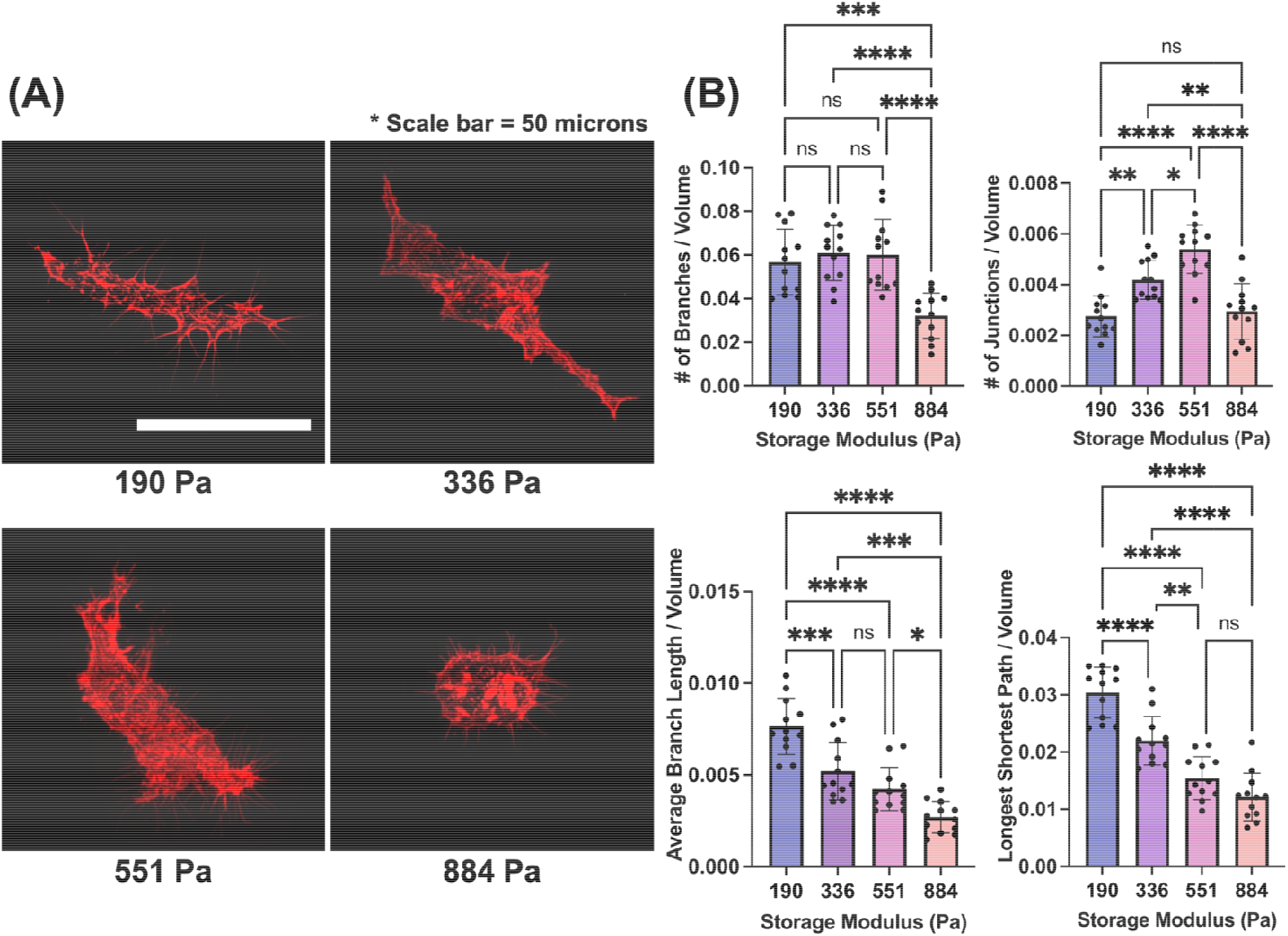
Hydrogel stiffness regulates cytoskeleton architecture in hiPSC-EPs following 4 days of culture. **(A)** Confocal images of F-actin (phalloidin, red) in hiPSC-EPs encapsulated in hydrogels of varying stiffness (190, 336, 551, and 884 Pa). Most extensive actin fibers are observed in cells encapsulated in 551 Pa hydrogels, while cells encapsulated in the 884 Pa hydrogel appear compact with reduced actin network. Scale bar = 50 µm. **(B)** Quantification of F-actin network complexity. Cells encapsulated in 551 Pa hydrogels had the highest branch and branching point numbers per unit volume, as well as larger average branch length and the longest shortest branch paths. Statistical significance is indicated as: *P ≤ 0.05, **P ≤ 0.01, ***P ≤ 0.001, ****P ≤ 0.0001

### 3.4 Matrix Stiffness Regulates Endothelial Differentiation in hiPSC-EPs

The dependence of endothelial differentiation on hydrogel stiffness was tracked by measuring the abundance of CD31, a critical marker of endothelial junction formation, in comparison to CD34, an endothelial progenitor marker. CD31/CD34 abundance ratio in hiPSC-EPs varied significantly depending on matrix stiffness, whereby cells encapsulated in the 551 Pa hydrogels exhibiting the highest ratio, nearly twice the CD31/CD34 ratios of the lower-stiffness hydrogels (190 Pa and 336 Pa) and 40% higher than the stiffer 884 Pa hydrogel (Fig. 4). The drop-off at 884 Pa likely reflects excessive stiffness disrupting cytoskeletal organization and reducing YAP/TAZ nuclear localization, which impairs mechanotransduction-driven CD31 expression while sustaining higher CD34 levels. Our results align with previously published data showing that increased matrix stiffness is associated with elevated CD31 expression in endothelial cells in both 2D and 3D culture systems [14, 16, 67]. This supports the role of mechanotransduction in regulating endothelial phenotype. Like cell morphology and the actin cytoskeleton, matrix stiffness of 551 Pa was optimal for endothelial differentiation.

**Figure 4:**
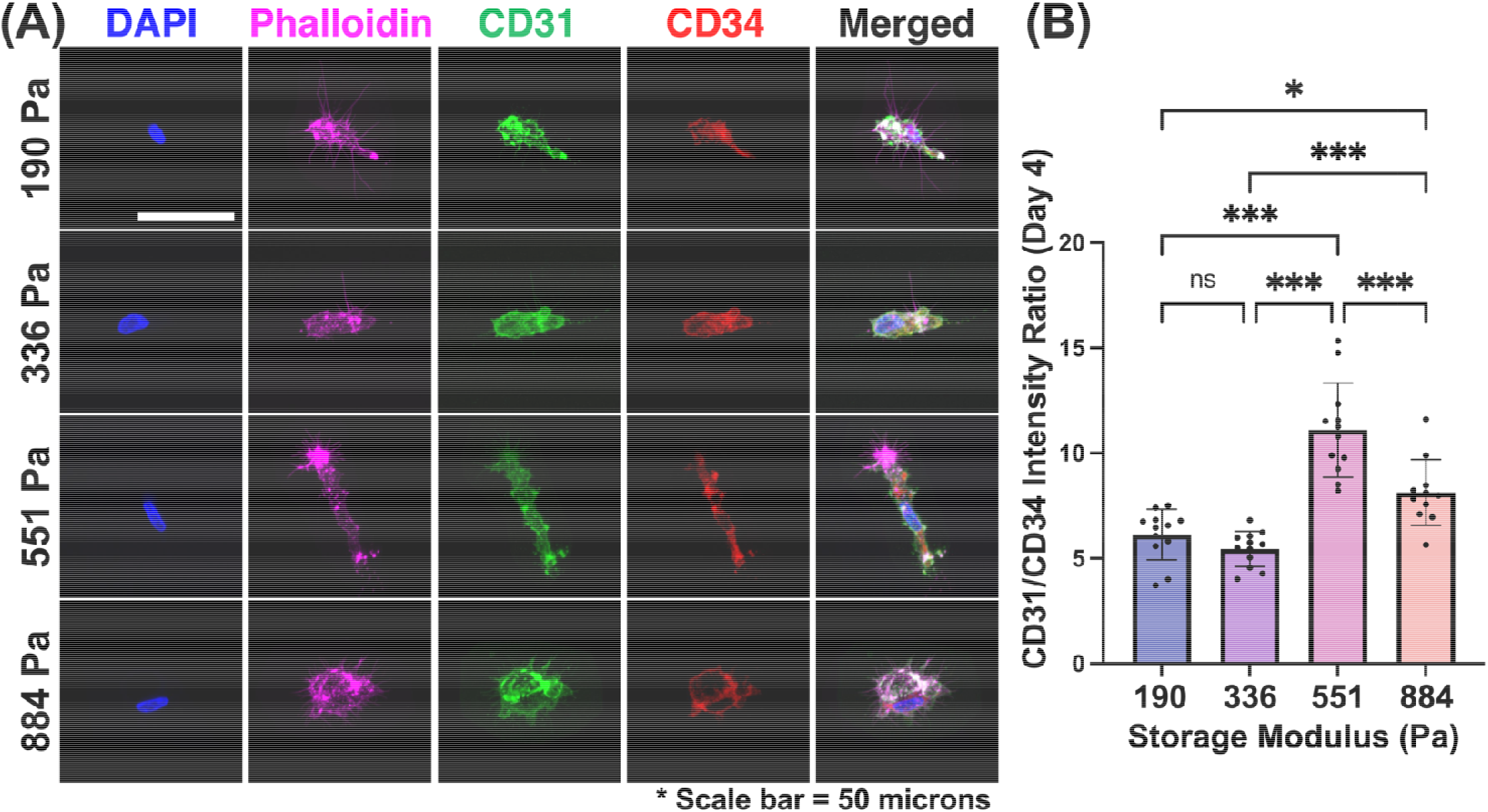
Abundance of endothelial markers in hiPSC-EPs cultured for 4 days in hydrogels of varying stiffness. **(A)** Representative immunofluorescence images showing nuclear staining (DAPI, blue), F-actin (phalloidin, magenta), CD31 (green), and CD34 (red). Scale bar = 50 microns. **(B)** Quantification of CD31/CD34 intensity ratio. Statistical significance is indicated as: *P ≤ 0.05, ***P ≤ 0.001.

### 3.5 NorHA hydrogel stiffness drives YAP nuclear expression but not TAZ

YAP and TAZ serve as primary mechanotransduction mediators in endothelial cells [40, 41], translocating between cytoplasm and nucleus in response to mechanical cues. YAP nuclear localization was analyzed in hiPSC-EPs as an indicator of mechanotransduction activity across different hydrogel stiffnesses. We hypothesized that the extent of mechanotransduction, measured as the ratio of YAP immunofluorescence signal in the nucleus compared to the cytoplasm will be stiffness-dependent, i.e. increase with the hydrogel stiffness in which the cells reside in. However, the nuclear-to-cytoplasmic YAP expression ratio peaked in cells encapsulated in the 551 Pa hydrogel, similar to the morphological responses. (Fig. 5).

**Figure 5:**
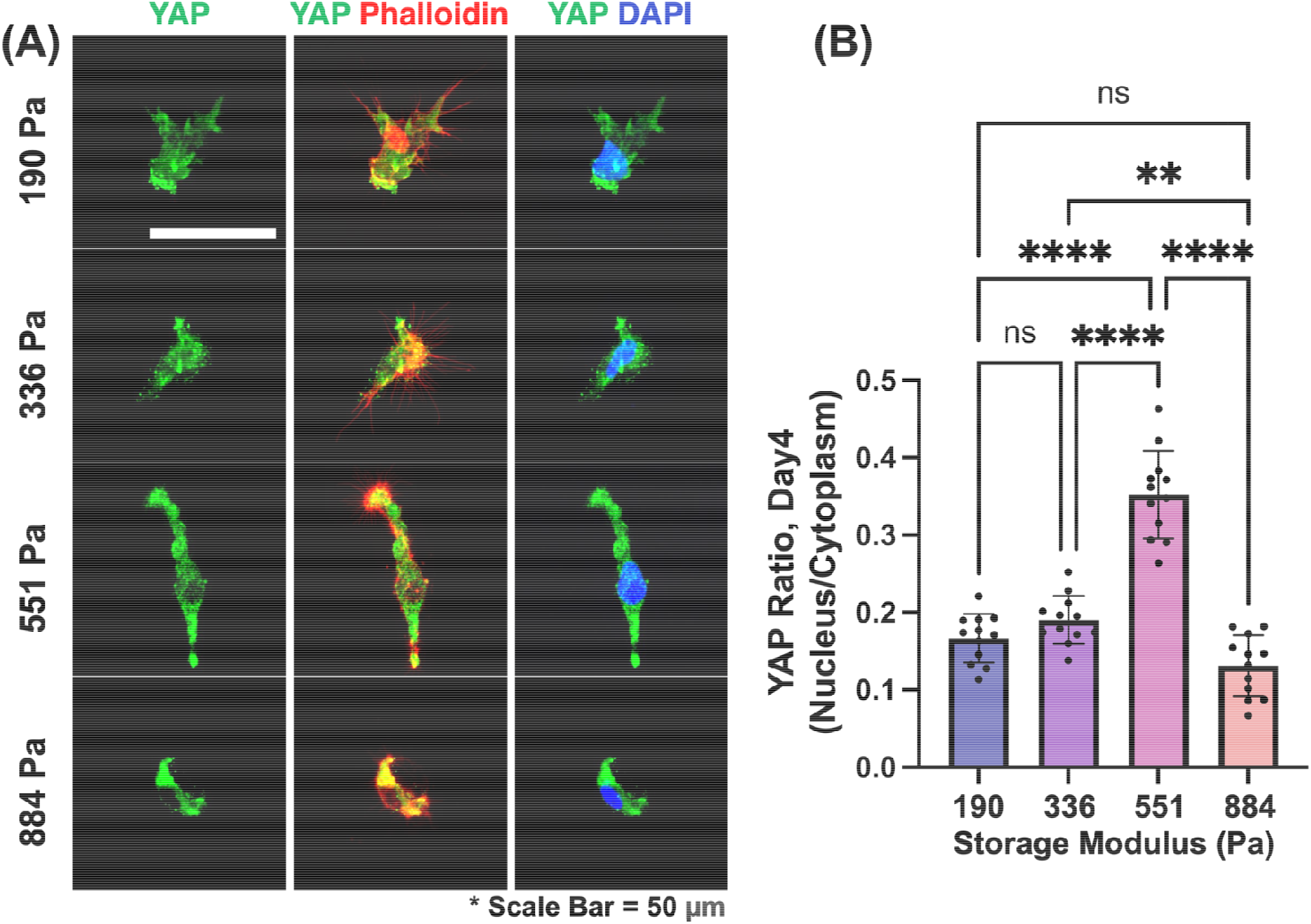
YAP localization analysis in hiPSC-EPs encapsulated in hydrogels of varying stiffnesses for 4 days. **(A)** Representative immunofluorescence images showing YAP (green), F-actin/phalloidin (red), and nuclear/DAPI (blue) expression in hiPSC-EPs encapsulated and cultured for 4 days in hydrogels with different storage moduli (scale bar = 50 µm). **(B)** Quantification of nuclear-to-cytoplasmic YAP ratio. hiPSC-EPs showed the highest nuclear localization when encapsulated in 551 Pa hydrogels. Statistical significance is indicated as: **P ≤ 0.01, ****P ≤ 0.0001.

To determine the changes in YAP abundance in each cellular compartment, the colocalization of the YAP signal was compared with either phalloidin, an F-actin marker, or DAPI, a nuclear marker. While YAP cytoplasmic abundance remained constant across all matrix stiffnesses, the nuclear abundance followed a similar pattern to the nucleus-to-cytoplasm YAP ratio (Supp Fig. 3). This suggests that the increased ratio resulted from nuclear translocation of a relatively minor cytoplasmic YAP fraction. Notably, nuclear YAP localization decreased by 54% in cells encapsulated in the 884 Pa hydrogels compared to those in softer matrices (190-551 Pa), indicating a mechanical threshold where increased matrix stiffness suppresses YAP nuclear translocation.

TAZ nuclear localization in hiPSC-ECs depicted a different response pattern from YAP when cultured in hydrogels of varying stiffness. While YAP exhibited peak nuclear localization in hiPSC-EPs encapsulated in 551 Pa hydrogels, the nuclear-to-cytoplasmic TAZ ratio in these cells remained consistently elevated across hydrogels with stiffness ranging from 190 to 551 Pa, showing no statistically significant differences among these conditions (Fig. 6). However, in hydrogels with a higher storage modulus (884 Pa), TAZ nuclear localization significantly decreased to a ratio of 0.05, representing approximately a 70% reduction compared to the lower-stiffness hydrogels (p < 0.0001). This reduction in both nuclear YAP and TAZ in hiPSC-Eps encapsulated in the 884 Pa hydrogels, suggests a mechanical threshold that influences mechanotransducer activity within physiologically relevant soft matrices.

**Figure 6:**
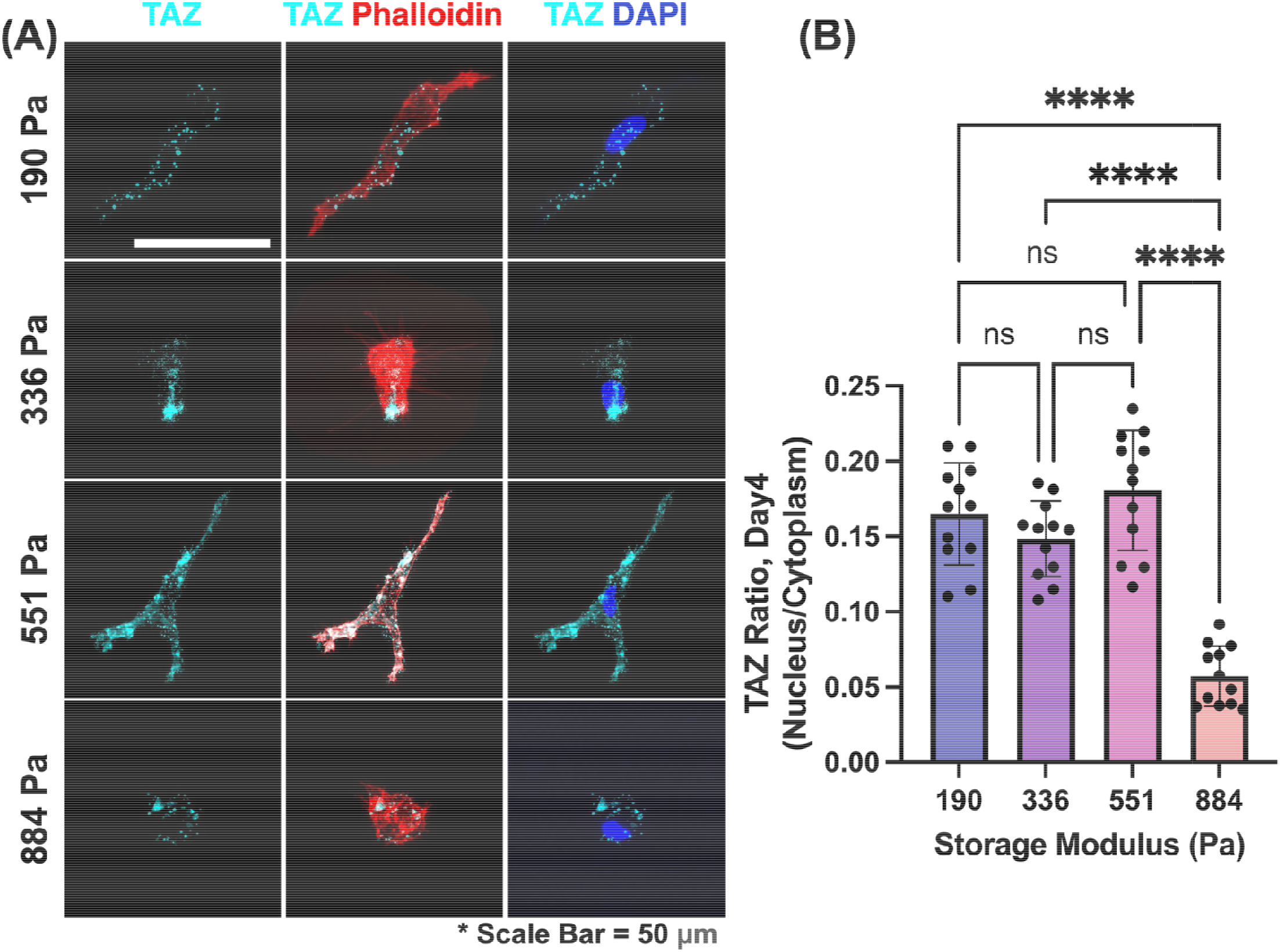
TAZ localization analysis in hiPSC-EPs encapsulated in hydrogels of varying stiffnesses for 4 days. **(A)** Quantification of nuclear-to-cytoplasmic TAZ ratio showing stable nuclear localization for cell encapsulated in hydrogels of 190-551 Pa with significant decrease when encapsulated in the stiffer 884 Pa hydrogels. **(B)** Representative immunofluorescence images showing TAZ (green), F-actin/phalloidin (red), and nuclear/DAPI (blue) staining across different storage moduli (scale bar = 50 µm). Statistical significance is indicated as: ****P ≤ 0.0001.

In 2D cell culture, nuclear YAP/TAZ localization increases with increasing substrate stiffness, for many cell types including HUVECs [20, 40]. However YAP/TAZ nuclear translocation in response to matrix stiffness in 3D environments remains less well understood, particularly in endothelial progenitors. For example, while YAP/TAZ nuclear localization in epithelial cells increased with increasing stiffness in 3D Matrigel/collagen matrices [68], for MSC cultured in alginate hydrogels [69], such a dependence was not observed. Similarly, Caliari et al. [70] observed that in HA hydrogels, YAP/TAZ nuclear localization in MSCs did not correlate with stiffness but instead depended on scaffold degradability. Our study reveals that a distinct biphasic res onse to stiffness occurs, peaking at 551 Pa before declining, while TAZ remains relatively stable across all conditions. This suggests that YAP is more sensitive to stiffness variations in 3D endothelial progenitors, whereas TAZ may be robustly regulated through alternative pathways.

### 3.6 YAP nuclear localization and CD31 abundance increased temporally in hiPSC-EPs

The analysis was then extended to day 7 to assess the temporal dynamics of mechanotransduction signaling. Representative immunofluorescence images demonstrate a marked shift in YAP distribution over time (Fig. 7A). At day 4, YAP signal (green) appeared diffuse throughout the cell with substantial cytoplasmic presence (Fig. 5), while by day 7, YAP showed pronounced nuclear enrichment, as evidenced by increased colocalization with DAPI staining (Supplementary Fig. 4). Overall, the trends observed in the day 4 cultures were also present in the day 7 cultures, with the highest nuclear YAP expression seen in the 551 Pa hydrogel (Fig. 7B). Notably, YAP expression significantly increased in the day 7 cultures across all tested stiffnesses, particularly in the 551 Pa condition. In this hydrogel stiffness, the nuclear-to-cytoplasmic YAP ratio increased by approximately 114% from day 4 (0.35 ± 0.05) to day 7 (0.75 ± 0.08) (Fig. 7B). This shift in YAP distribution was further validated through Pearson’s correlation analysis (Supplementary Fig. 3). YAP-nuclear colocalization increased by 95% from day 4 to day 7 (0.38 ± 0.03 to 0.74 ± 0.05), while YAP-cytoplasmic colocalization decreased by 32% (0.95 ± 0.03 to 0.65 ± 0.05). Similarly, CD31/CD34 ratio increased across all stiffness conditions in the 7-day cultures, with the highest ratio at the 551 Pa hydrogel, exhibiting approximately 150% increase from day 4 (10.2 ± 1.5) to day 7 (25.4 ± 2.1) (Fig. 8). This temporal increase in the CD31/CD34 ratio indicates that endothelial differentiation proceeded and was still ongoing on day 7 of culture. In the 7-day cultures, we observed a stiffness-dependent actin cytoskeleton architecture. Specifically, the number of branches increased with stiffness by approximately 46% from the softest (190 Pa, 0.041±0.013) to the stiffest hydrogel (551 Pa, 0.060±0.012). Conversely, other cytoskeletal parameters decreased with increasing stiffness: the number of junctions decreased by 25% (from 0.008±0.002 at 190 Pa to 0.006±0.002 at 551 Pa), average branch length decreased by 68% (from 0.031±0.006 μm/μm^3^ at 190 Pa to 0.010±0.004 μm/μm^3^ at 551 Pa), and longest shortest path decreased by 69% (from 0.083±0.034 at 190 Pa to 0.026±0.012 at 551 Pa) (Supplementary Fig. 5). This suggests that in stiffer hydrogels, endothelial progenitors generate more actin protrusions, likely as an adaptive response to mechanical resistance. However, these branches remain less interconnected, forming longer, more structured fibers rather than a dynamically connected network. This shift may indicate that cytoskeletal reinforcement takes precedence over reorganization, potentially driven by higher YAP/TAZ activity, which promotes actin stress fiber elongation rather than de novo branching. Over 7 days of culture, the actin cytoskeleton network became more developed compared to day 4, as reflected by substantial increases in cytoskeletal parameters across all hydrogel stiffness levels. The average branch length increased by approximately 1.24-fold (from ∼0.025 to 0.031±0.006 μm/μm^3^) at 190 Pa, 1.6-fold (from ∼0.01 to 0.016±0.005 μm/μm^3^) at 336 Pa, and 2-fold (from ∼0.005 to 0.010±0.004 μm/μm^3^) at 551 Pa. Similarly, the longest shortest path increased by 1.66-fold (from ∼0.05 to 0.083±0.034) at 190 Pa, 1.67-fold (from ∼0.03 to 0.050±0.027) at 336 Pa, and 1.3-fold (from ∼0.02 to 0.026±0.012) at 551 Pa) across all hydrogel stiffness levels. However, the specific patterns of actin organization differed based on substrate stiffness. In stiff hydrogels, the number of branches and junctions remained unchanged between day 4 and day 7. Instead of forming additional connections, cells reinforced existing structures, elongating their cytoskeletal fibers to accommodate mechanical stress. This stabilization suggests a plateau in cytoskeletal remodeling over time. In contrast, compliant hydrogels (190 Pa) exhibited a 45% increase in junctions over time (from ∼0.0055 at day 4 to 0.008±0.002 at day 7), indicating a transition toward a more interconnected actin cytoskeleton. This suggests that softer environments permit sustained cytoskeletal remodeling, leading to greater interconnectivity and adaptability. Following the observations in the 880 Pa hydrogel, where cells exhibit a rounded morphology and display the lowest CD31 and YAP expression at both 4 and 7 days, we inferred that the cells are not well-spread or fully engaged with the matrix, and their mechanosensitive pathways are minimally activated. Consequently, we did not study the actin architecture in these hydrogels, as they are less informative regarding cytoskeletal dynamics and force generation.

**Figure 7:**
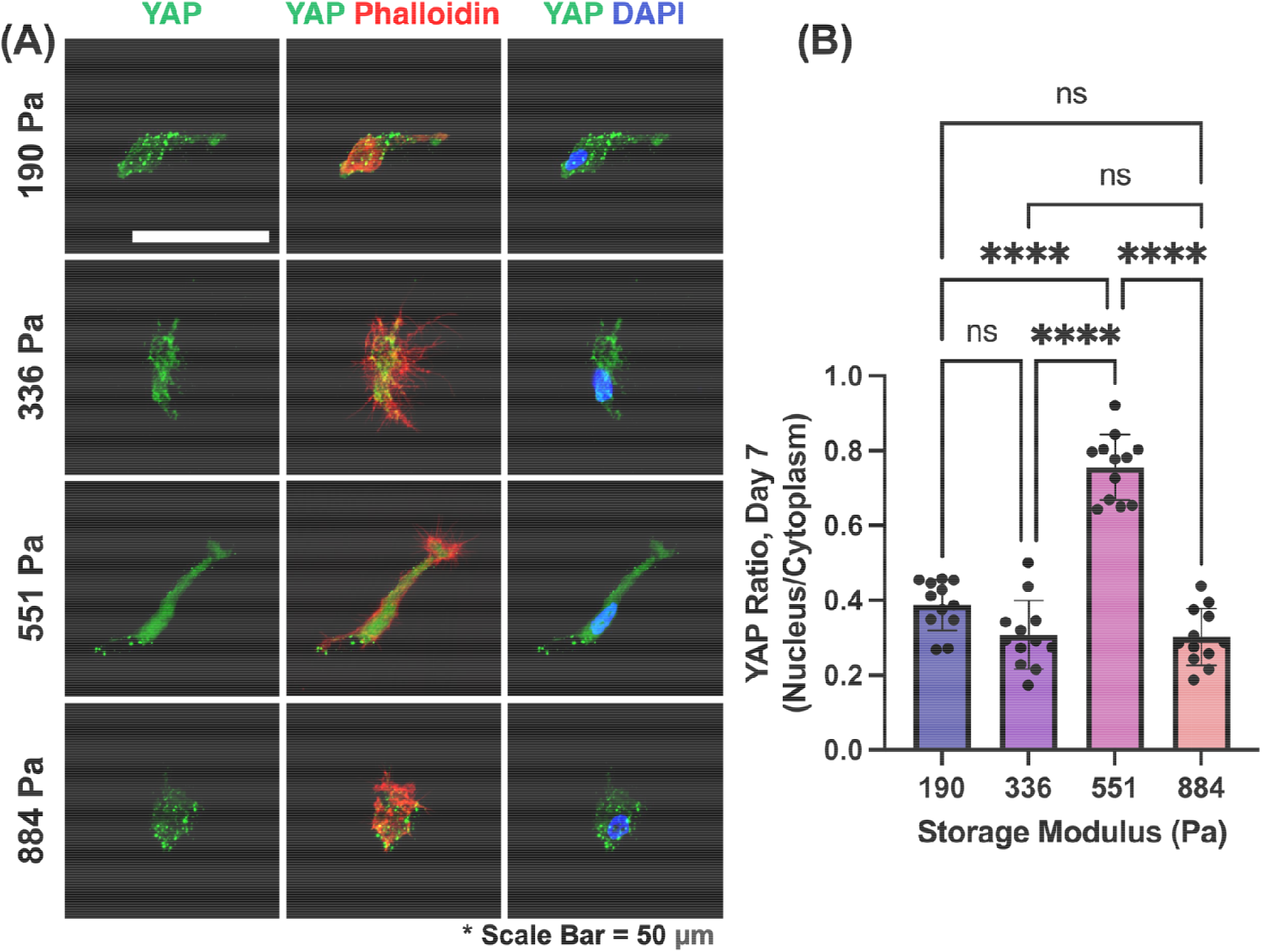
Effect of culture duration on YAP localization in hiPSC-EPs. **(A)** Representative immunofluorescence images showing YAP (green), F-actin/phalloidin (red), and nuclear/DAPI (blue) staining of hiPSC-EPs encapsulated in hydrogels of varying stiffness. Scale bar = 50 μm. **(B)** Nuclear-to-cytoplasmic YAP localization ratio had comparable levels in cells encapsulated in 190 Pa and 884 Pa hydrogels, bue was significantly higher in cells encapsulated in the 551 Pa hydrogels. Statistical significance is indicated as: ****P ≤ 0.0001; ns = not significant.

**Figure 8:**
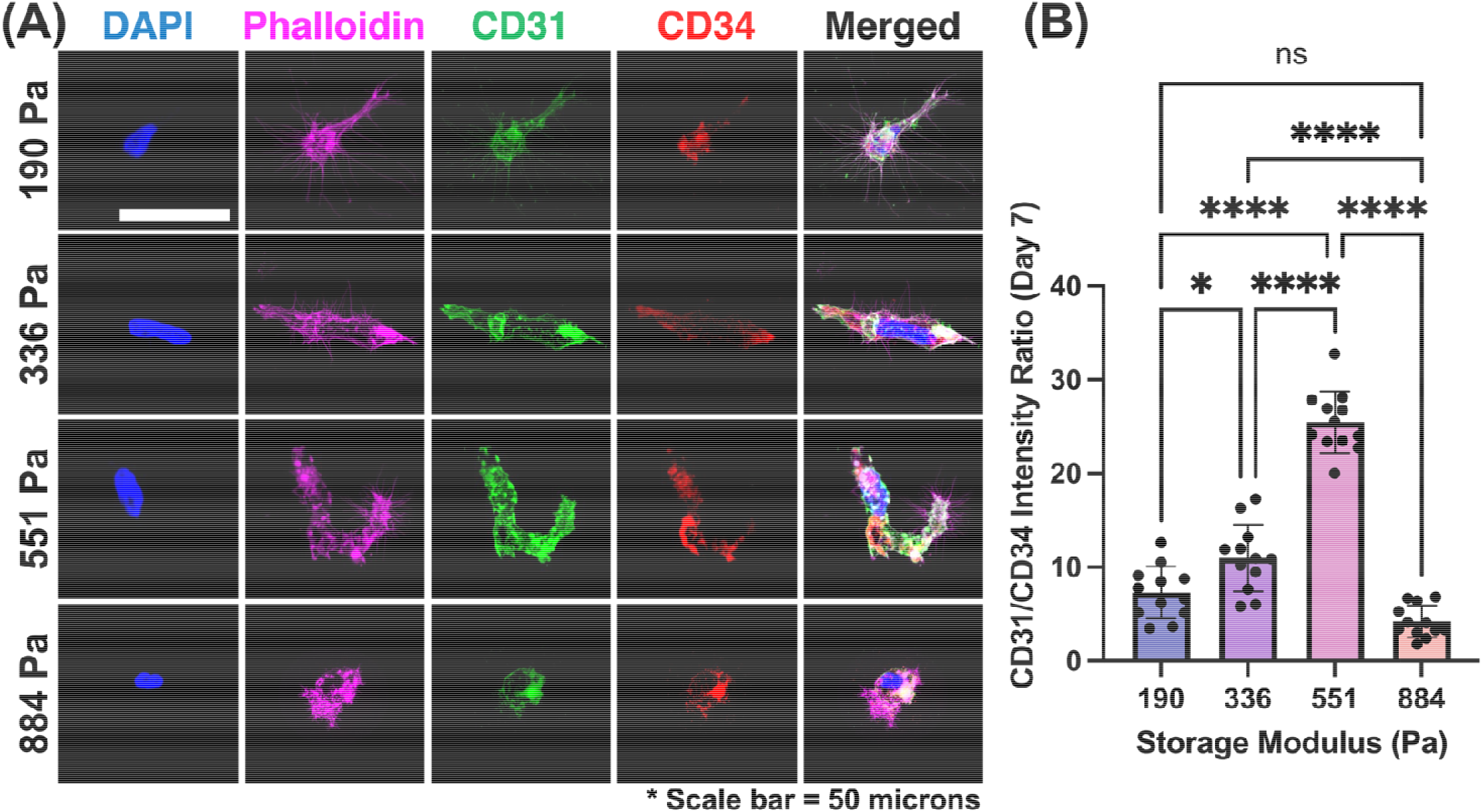
Abundance of endothelial markers in hiPSC-EPs encapsulated in hydrogels of varying stiffness for 7 days. **(A)** Representative immunofluorescence images showing nuclear/DAPI (blue), F-actin/phalloidin (magenta), CD31 (green), and CD34 (red) staining across different storage moduli (scale bar = 50 μm). **(B)** Quantification of CD31/CD34 abundance ratio at day 7 showing progressive increase as cells are encapsulated in hydrogels of increasing stiffness from 190 Pa to 551 Pa and a significant decrease at 884 Pa. Statistical significance is indicated as: *P ≤ 0.05, ****P ≤ 0.0001; ns = not significant.

### 3.7 3D basal contractility of hiPSC-EPs depends on culture maturity and hydrogel compliance

To determine how hydrogel stiffness and culture duration affect the basal contractility of hiPSC-EPs, 3D traction force microscopy was employed to measure the cellular displacements. The actin polymerization inhibitor, cytochalasin D (CytoD), was employed to cause full relaxation of the cells. To determine the basal contractility of the cells, the data was analyzed from the CytoD-treated state to the basal state. To track the cell’s movement in the hydrogel, the hydrogel is impregnated with fluorescent bead fiducial markers which are imaged before and after encapsulated hiPSC-EPs were treated with CytoD [71]. Production of a Gaussian process (GP) model from the bead displacements allows for the determination of displacements along the finite element mesh representation of the cell (Fig 9. A-B).

**Figure 9:**
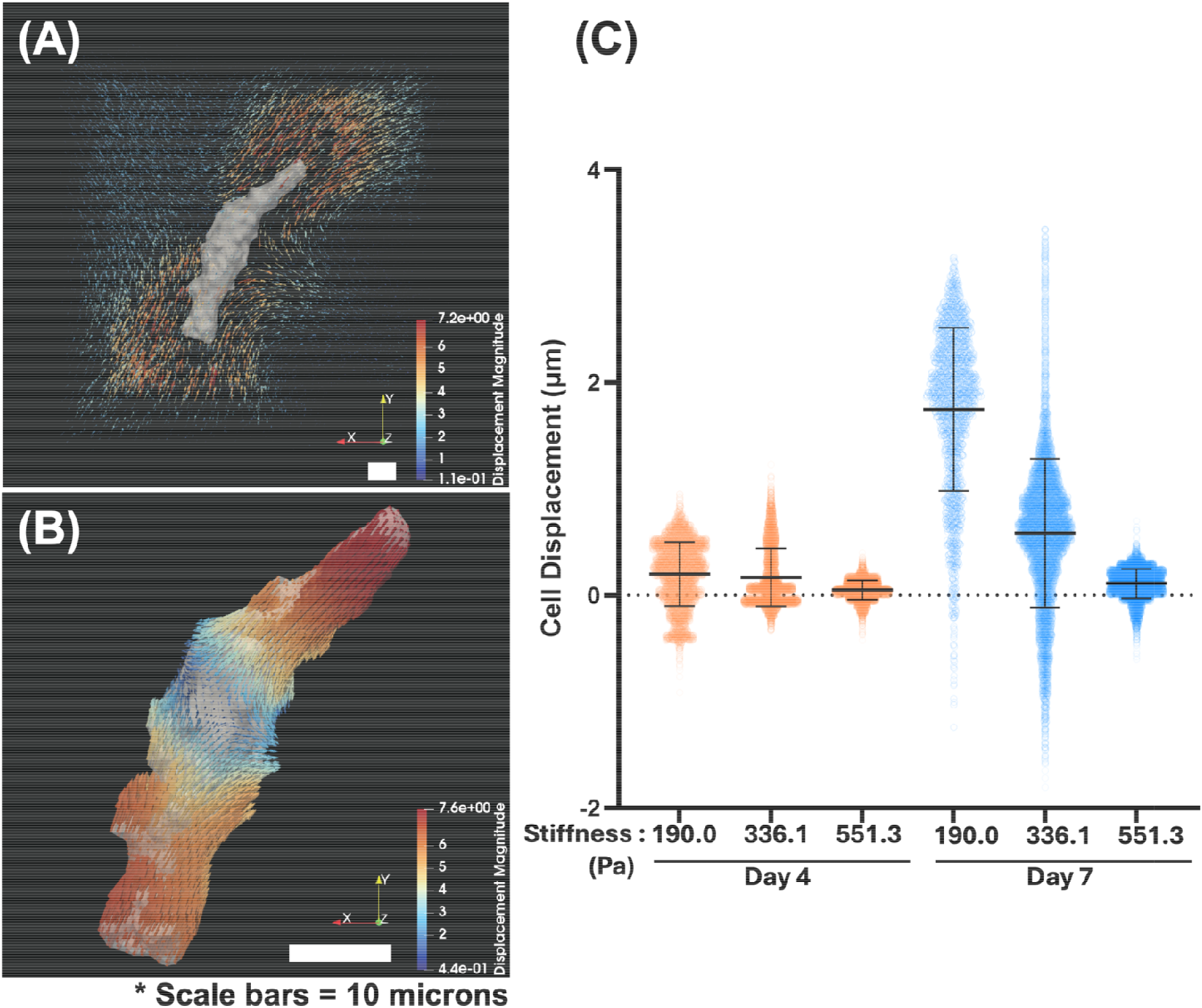
hiPSC-EP basal contractility depends on culture maturity and hydrogel stiffness. **(A)** hydrogel displacements are determined by nearest neighbor tracking and filtering of fluorescent microspheres in the gel, which are then input to produce a Gaussian process (GP) model. Scale bar = 10 µm. **(B)** The GP model is analyzed at the points of the cell surface mesh to determine cell surface displacements. Scale bar = 10 µm. **(C)** hiPSC-EP surface displacements measured from the relaxed, cytochalasin D-treated state to the basal state. Displacements are presented with respect to the negative of the dot product of the normal to the cell surface. Mean ± SD plotted as lines on top of an all-data point spread. All groups are significantly different from each other (p<0.0001) by two-way ANOVA and Tukey post-test.

hiPSC-EPs were imaged following encapsulation in hydrogels of average stiffness of 190, 336, or 551 Pa after 4 or 7 days in culture (Fig 9C). As stiffness increases, overall displacement caused by basal contraction significantly decreases nonlinearly (), with day 4 displacements decreasing from 0.2008 ± 0.3015 μm in 190 Pa gels to 0.05164 ± 0.08992 μm in 551 Pa gels. Similarly, basal contractility after 7 days of culture was found to also nonlinearly decrease with hydrogel stiffness (simple linear regression R2=0.35), with day 7 displacements decreasing from 1.748 ± 0.7658 μm in 190 Pa gels to 0.1120 ± 0.1378 μm in 551 Pa gels. These results depict that hydrogel stiffness greatly influenced basal cell displacements, with stiffer hydrogels causing less displacement of the hiPSC-EPs in a nonlinear manner. Moreover, the displacements of these cells portray that increasing maturity of the culture greatly increases the basal contractility of the hiPSC-EPs. Few studies have applied 3D traction force microscopy (3D TFM) to endothelial cells’ response to matrix stiffness. Shapeti et al. [72] examined HUVECs in 3D PEG hydrogels (433–667 Pa), comparing CCM2-silenced (mutant) and wild-type cells. Loss of CCM2, a key regulator of RhoA/ROCK signaling and cytoskeletal organization, increased actomyosin contractility and traction forces, making cells more sensitive to stiffness. In stiffer environments, this heightened mechanical stress contributed to disrupted junctions and pathological vascular remodeling, mirroring cerebral cavernous malformations. Ucla et al. [73] introduced a method to quantify

HUVEC traction forces at the fiber scale using deformable PEGDA-collagen arrays (3.5–17.6 Pa). Their findings, aligning with ours, showed that softer matrices permitted larger displacements due to lower mechanical resistance, while stiffer matrices limited displacement but led to increased traction forces as cells exerted more force to deform the rigid environment.

### 3.8 hiPSC generate the most developed microvascular plexus in complaint hydrogels

Lastly, we evaluated how hydrogel stiffness impacts the ability of hiPSC-EPs’ plexus format on and connectivity. Interconnected, quantifiable vascular networks formed in hydrogels across the tested stiffness range. Using our computational pipeline [26, 74], we analyzed the microvascular networks’ topology (i.e., length and connectivity). The vasculogenic potential of each hydrogel was quantified by calculating the number of branch and endpoints points (both indicators of network complexity), network volume fraction (which represents the total volume occupied by vascular structures within the hydrogel), and network connectivity (a measure of the frequency of vessel junctions.

Overall, our findings indicate that hiPSC-EPs generated the most extensive and well-connected vascular networks in the most compliant hydrogel condition, with a stiffness of 190 Pa, aligning with our previously published data using an interpenetrating polymer network of collagen and NorHA [29]. As the hydrogel stiffness increased, the complexity and connectivity of the vascular networks progressively declined. Notably, the most limited network topology was observed in the stiffest hydrogel tested, with a stiffness of 880 Pa (Fig. 10A). Quantitative analysis revealed that compliant hydrogels (190 Pa) supported 28.5-fold higher branch formation (Fig. 10B), 2.3-fold more endpoints (Fig. 10C), 4.9-fold greater network volume fraction (Fig. 10D), and 3-fold higher connectivity (Fig. 10E and Supplementary Video 1) compared to the stiffest condition (884 Pa). These results clearly demonstrate the significant impact of matrix stiffness on vascular network development, with the most extensive network formation occurring in more compliant environments.

**Figure 10:**
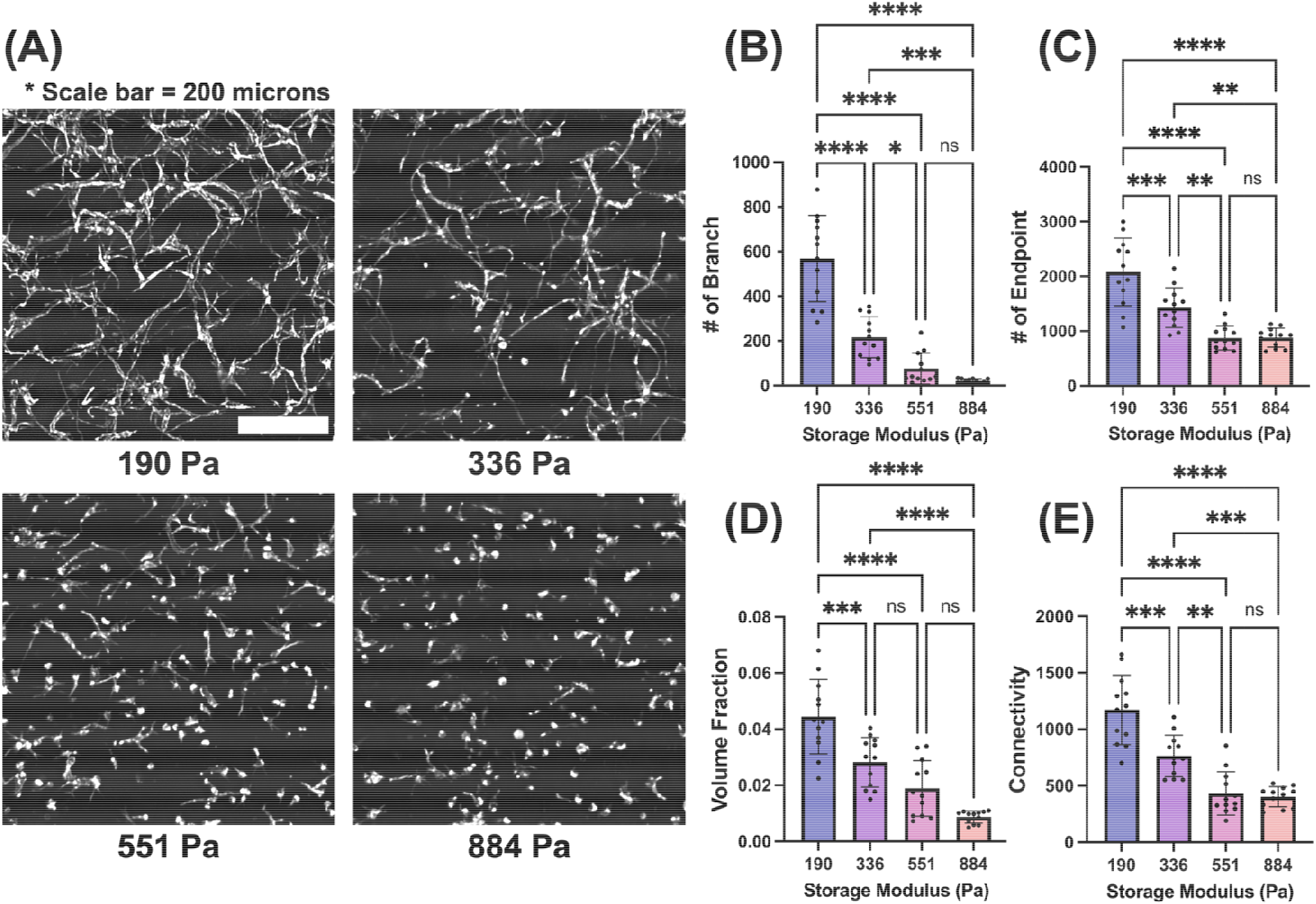
Hydrogel stiffness influences the vasculogenic potential of hiPSC-EP. **(A)** Representative images of hiPSC-EP-derived microvascular networks formed within hydrogels of varying storage moduli (190, 336, 551, and 884 Pa) following 7 days of culture. Scale bar = 300 µm. **(B)** Quantification of branch points showing progressive decrease with increasing hydrogel stiffness. **(C)** Number of endpoints decreases significantly as hydrogel stiffness increases. **(D)** Volume Fraction measurements show the 190 Pa hydrogel supports the highest vascular volume, which progressively decreases with increasing stiffness. (E) Connectivity decreases significantly in stiffer matrices. Statistical significance indicated as: (*p<0.05, **p<0.01, ***p<0.001, ****p<0.0001) by one-way ANOVA and Tukey post-test.

## 4. Discussion

In this study, we aimed to investigate how matrix stiffness regulates endothelial cell differentiation and mechanotransduction using norbornene-modified hyaluronic acid hydrogels with precisely controlled mechanical properties. We developed a unique ECM-mimicking system that allows modulation of stiffness without altering the hydrogel’s chemical components. or degradation profile. This system enabled a systematic investigation of how hydrogel stiffness influences cell morphology, differentiation markers, and mechanotransduction events. Notably, the stiffness ranges employed in this study fall within the physiological spectrum of vascular development [7-9], making these findings highly relevant to *in vivo* conditions.

The current results demonstrate that hydrogel stiffness is a key regulator of endothelial cell morphology, mechanosensitive signaling, adhesion protein expression, and cytoskeletal contractility. Moreover, there is a pronounced dependence of endothelial cell morphology—measured by volume and surface area—on hydrogel stiffness. Cells cultured in intermediate-stiffness hydrogels (551 Pa) exhibited the highest volume and surface area, reflecting their most spread-out morphology with numerous protrusions. In contrast, cells in lower-stiffness hydrogels remained extended but had smaller volumes and surface areas, while those in higher-stiffness hydrogels were rounded, exhibiting the lowest volume and surface area. This behavior is reflected by the actin filament network architecture of these cells. Specifically, we noted a transition from single, elongated actin filaments to an extensively interconnected network, followed by a shift to fragmented and disorganized filaments at higher stiffness levels. The impact of stiffness on cell morphology, particularly in endothelial cells, has been well studied in both 2D and 3D microenvironments, demonstrating a general trend of increased cell spreading with increasing stiffness [22, 75]. However, most 2D studies utilize stiffness ranges that are significantly higher than the physiological endothelial microenvironment, making them more representative of pathophysiological conditions. Conversely, while many 3D studies are more relevant to physiological stiffness, they often involve systems where multiple parameters, such as available cell-binding ligands, or degradability, are varied alongside stiffness, making it difficult to isolate stiffness specific effects. This underscores the significance of our fully 3D approach to quantify cell contraction patterns and subsequent findings, highlighting hydrogel stiffness as a critical determinant of cell morphology and behavior.

Notably, we found that both CD31 expression and YAP nuclear translocation exhibited a similar dependence on hydrogel stiffness, reaching their highest levels at intermediate stiffness. This potentially suggests that their regulation is linked through cytoskeletal tension and mechanotransduction pathways, where highly branched interconnected actin filament organization supports both YAP activation and CD31 expression, while excessive stiffness disrupts this balance. Although CD31 is primarily associated with endothelial cell-cell adhesion and junction maintenance, its increased expression in iPSC-EP encapsulating in hydrogels of increasing stiffness suggests additional roles, potentially in cytoskeletal anchoring and mechanosensitive signaling [75-78]. YAP, a key mechanotransducer, is essential for translating substrate stiffness and cell shape into biochemical signals. YAP activity is regulated by several cell surface receptors of extracellular matrix proteins, including collagen, fibronectin, vitronectin, and thrombospondin [79, 80]. Importantly, YAP activation has been implicated in angiogenesis, as recently reviewed by Azad et al. [82] and is further influenced by vascular endothelial growth factor (VEGF), which promotes YAP/TAZ activity via its impact on the actin cytoskeleton [45]. These findings collectively reinforce the concept that matrix stiffness modulates YAP activation, which in turn increases CD31 expression in endothelial cells. Nonetheless, it is important to acknowledge that CD31 expression may also be regulated by additional signaling pathways beyond YAP alone. Interestingly, TAZ nuclear expression did not follow the same trend as YAP; instead, it remained unchanged across the studied stiffness range and decreased at the highest stiffness. This suggests that the two transcription factors have differential roles in vasculogenesis across the studied stiffness [81]. Notably, TAZ has been reported to be more sensitive to cell-cell junctions, whereas YAP is more influenced by substrate stiffness and cell spreading [41, 83]. The relationship between CD31 expression and YAP nuclear translocation with hydrogel stiffness further underscores the role of mechanosensing in endothelial cell function. It also aligns with the concept that endothelial cells sense and respond to their mechanical environment, which in turn modulates their morphology, adhesion, and function. To further explore how these mechanosensitive pathways influence cellular mechanics, we examined the extent of cell basal contractility by measuring displacement following CytoD treatment, which provides insight into the extent of actomyosin-driven tension and the mechanical resilience of the cytoskeletal network. Displacement following CytoD treatment decreases nonlinearly with increasing hydrogel stiffness, with cells in stiffer hydrogels exhibiting less displacement. In compliant matrices (190 Pa), the actin filament network is more interconnected with longer branches and a greater longest shortest path, allowing for more dynamic cytoskeletal reorganization. When actin is disrupted, these cells experience greater relaxation and spreading, resulting in higher displacement. In stiffer matrices (551 Pa), the actin filament network has more junctions but shorter branches and a smaller longest shortest path, forming a more compact, crosslinked cytoskeleton. This denser structure limits large-scale relaxation, preventing significant cytoskeletal collapse when CytoD is applied, leading to lower displacement. Instead of relying on long-range actin connectivity, cells in stiff matrices stabilize their shape through localized actin reinforcement, restricting their response to actin depolymerization.

An important observation emerged when comparing 4-day and 7-day cultures. At 7 days, cells exhibited higher CD31 expression, greater YAP nuclear translocation, and larger displacement following CytoD treatment, indicating increased cytoskeletal maturation and contractile response over time. This trend was observed across all hydrogel stiffness levels except for the 551 Pa hydrogel, where displacement remained unchanged between the two time points. This suggests that over time, cells develop more interconnected and contractile cytoskeletal networks, which store greater mechanical tension that is released upon actin disruption, leading to increased displacement. In compliant hydrogels, where junction density increased over time, this enhanced interconnectivity likely facilitated force transmission and relaxation upon CytoD treatment. However, in the stiffer hydrogels, while cytoskeletal fibers became longer, the number of branches and junctions remained stable, suggesting that cells reached a mechanically stable configuration. This structural reinforcement may have counteracted further deformation, limiting displacement despite increased intracellular tension. These findings underscore the interplay between intracellular contractility and extracellular matrix mechanics: in compliant environments, prolonged culture allows for greater cytoskeletal remodeling and tension buildup, enhancing displacement upon actin disruption, whereas in stiffer environments, mechanical stabilization of the actin filament network prevents further deformation, overriding the cells’ ability to contract and deform.

Previously, we observed that in systems where cells fail to elongate by Day 4, interconnected vascular networks do not form [29, 59]. To further investigate the mechanostructive properties of this hydrogel system, we conducted this study to understand how matrix mechanics influence endothelial behavior and network formation. Indeed, in the stiffest hydrogel tested (880 Pa), limited cell elongation and reduced network connectivity were observed, consistent with our earlier findings. However, for the other hydrogel conditions, cellular displacement emerged as a key determinant of network formation. Specifically, while the 551 Pa hydrogel exhibited the highest surface volume, area, and expression of nuclear YAP as well as CD31/CD34, it did not support the most interconnected and robust microvascular network. Instead, the most compliant hydrogel (190 Pa) facilitated superior network formation. This hydrogel condition also demonstrated the highest displacement values following CytoD treatment, underscoring that cellular displacement within the matrix is a critical driver of microvascular organization. Our findings provide valuable insights into the dynamic interplay between mechanical cues and cellular responses, which could inform the design of better biomaterials for vascular tissue engineering and related applications.

### Conclusion and future implications

The results from this study have significant implications for the development of vascular and angiogenic hydrogels, particularly for tissue engineering and regenerative medicine applications. The ability to control endothelial cell behavior through hydrogel stiffness is critical for designing biomaterials that can promote vascularization or support the growth of functional blood vessels in engineered tissues. By optimizing the mechanical properties of hydrogels, angiogenesis and potentially the integration of engineered tissues with the host vasculature could be better supported. However, hydrogels that are too stiff may hinder endothelial cell migration and vascular sprouting, processes that are necessary for effective angiogenesis, due to excessive matrix resistance. Therefore, for angiogenic hydrogel applications, a careful balance of matrix stiffness is required to promote cell migration, network formation, and blood vessel maturation while preventing excessive mechanical restriction. Additionally, the finding that displacement increases with time in culture (at least in lower to intermediate stiffness hydrogels) suggests that long-term culture may be necessary for endothelial cells to mature sufficiently to promote vascular sprouting and network formation. This insight will be particularly useful when developing hydrogels for applications such as wound healing, vascular grafts, and organ regeneration, where long-term tissue development is required for the formation of stable, functional blood vessels. Incorporating time-dependent factors into hydrogel design, in addition to optimizing stiffness, will help engineers create dynamic environments that support endothelial cell maturation and angiogenic potential over extended periods. This will be crucial for ensuring that the engineered vascular structures remain stable and functional over time as they integrate with host tissues. A limitation of this study is that stiffness and mesh size are coupled in the 3D hydrogels, which may impact biological signaling in addition to mechanosignaling. Future work could leverage strategies to independently tune the stiffness of the hydrogel, e.g., through the structure of the macromer [84, 85] or crosslinkers [86]. Ultimately, by fine-tuning hydrogel mechanical properties, considering cellular maturation dynamics, and optimizing matrix interactions, more effective angiogenic materials that enhance vascularization in tissue engineering and regenerative medicine can be developed. In addition, we are conducting inverse computational modeling to determine cell stress fiber behavior, as in [87]. These advances will bring us closer to engineering vascularized tissue constructs, organ regeneration platforms, and transplantation models that rely on robust and functional vasculature to sustain long-term viability.

## Acknowledgments

We gratefully acknowledge the support of the National Institutes of Health (R01HL15829 awarded to J.Z., R35GM138193 awarded to A.M.R, F31 HL170717-02, 5F32HL167570 awarded to T.W.) and the National Science Foundation (MRSEC DMR-2308817 awarded to A.M.R.). J.Z. and J.H would like to thank Dr. Arie Horowitz for insightful discussions.

## Author Contributions

J.H: Data curation, Formal analysis, Investigation, Visualization, Software, Writing – original draft, Writing – review and editing. K.H.: Data curation, Formal analysis, Investigation, Visualization, Writing – original draft, Writing – review and editing. T.W.: Data curation, Formal analysis, Investigation, Visualization, Software, Writing – original draft, Writing – review and editing. B.L.: Data curation, Writing – review and editing. M.S.S.: Funding acquisition, Resources, Supervision, Formal analysis, Writing – review and editing. A.M.R.: Conceptualization, Funding acquisition, Resources, Project administration, Supervision, Writing – original draft, Writing – review and editing. J.Z.: Conceptualization, Funding acquisition, Resources, Project administration, Supervision, Writing – original draft, Writing – review and editing.

## Author Declarations

No interests are present to disclose.

## Data Availability

The data that support the findings of this study are available from the corresponding author upon reasonable request.

