## Supplementary Fig for "Matrix Stiffness Regulates Mechanotransduction and Vascular Network Formation of hiPSC-Derived Endothelial Progenitors Encapsulated in 3D Hydrogels"

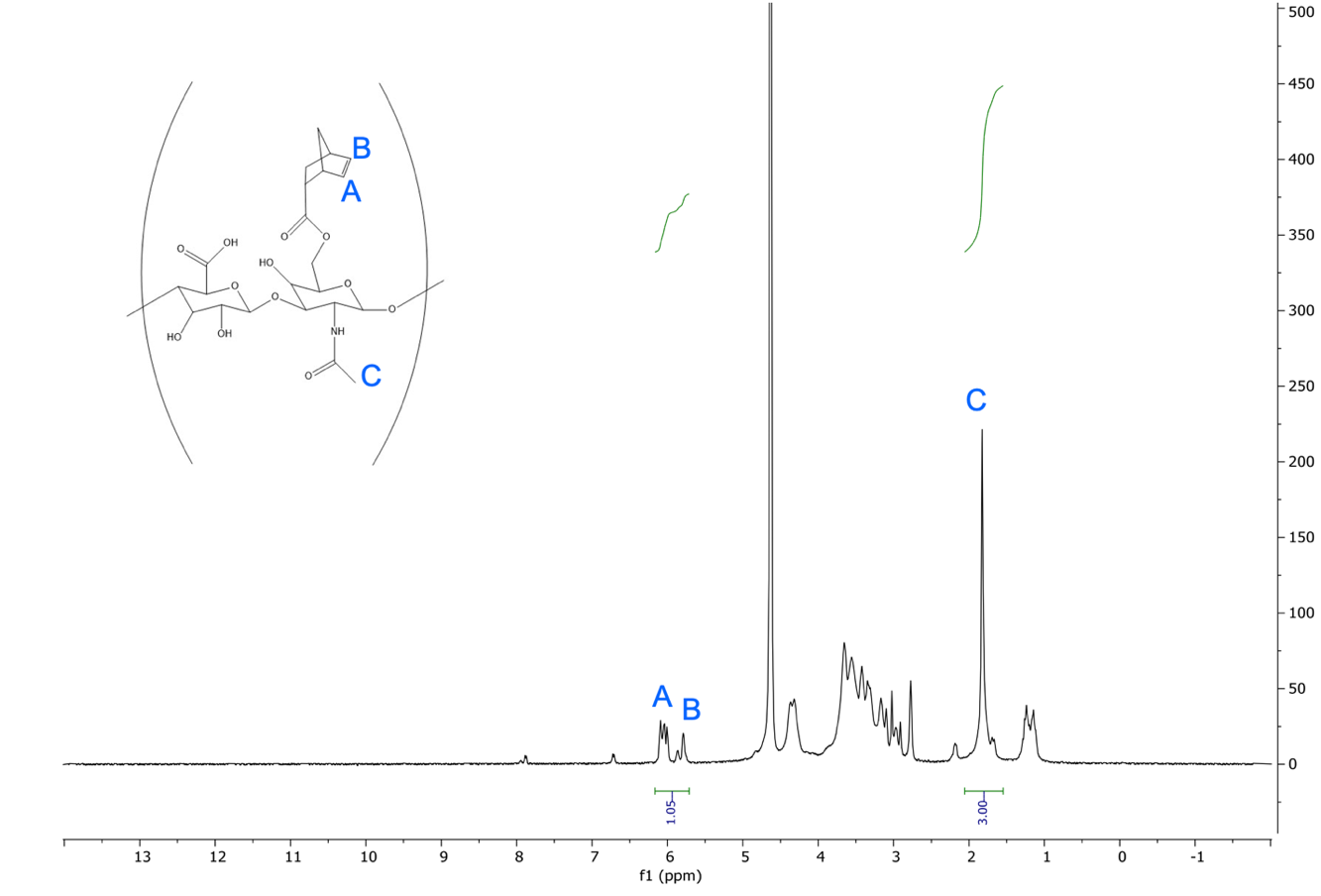

**Supplementary Figure 1:** ¹H NMR Characterization of NorHA Functionalization in D₂O. ^1^H NMR spectrum, confirming NorHA functionalization with norbornene at approximately 52.5%. Characteristic norbornene peaks were integrated relative to the HA backbone to calculate the extent of modification.

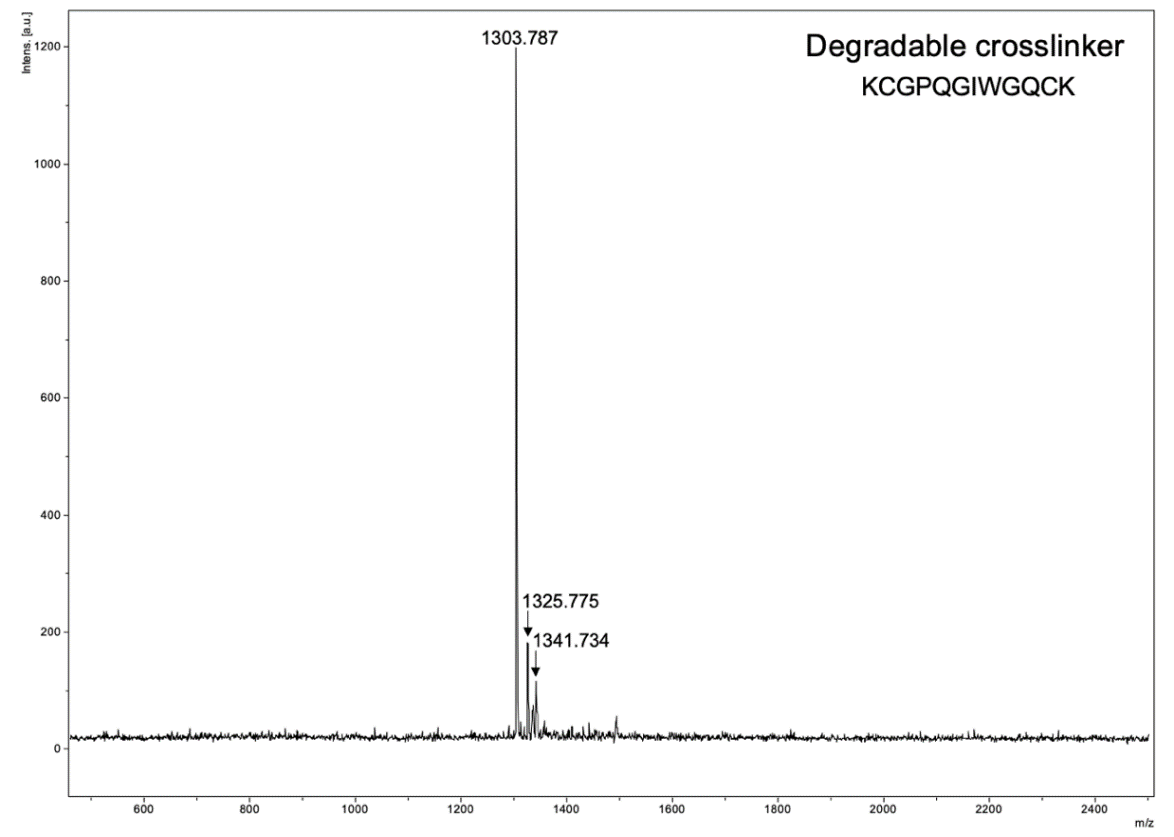

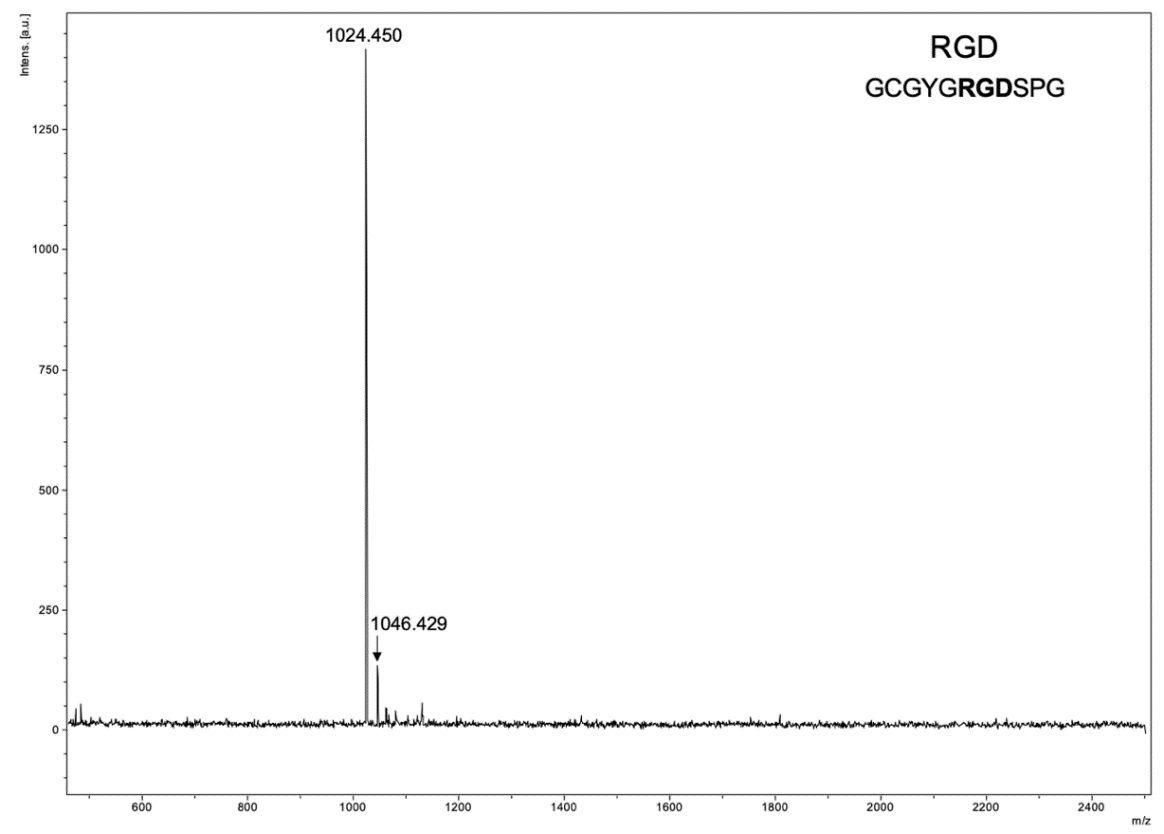

**Supplementary Figure 2:** Matrix-Assisted Laser Desorption/Ionization (MALDI) characterization of degradable peptide crosslinker and RGD Motif. MALDI spectra confirm the expected molecular weights of both the peptide crosslinker and the RGD cell-adhesive sequence.

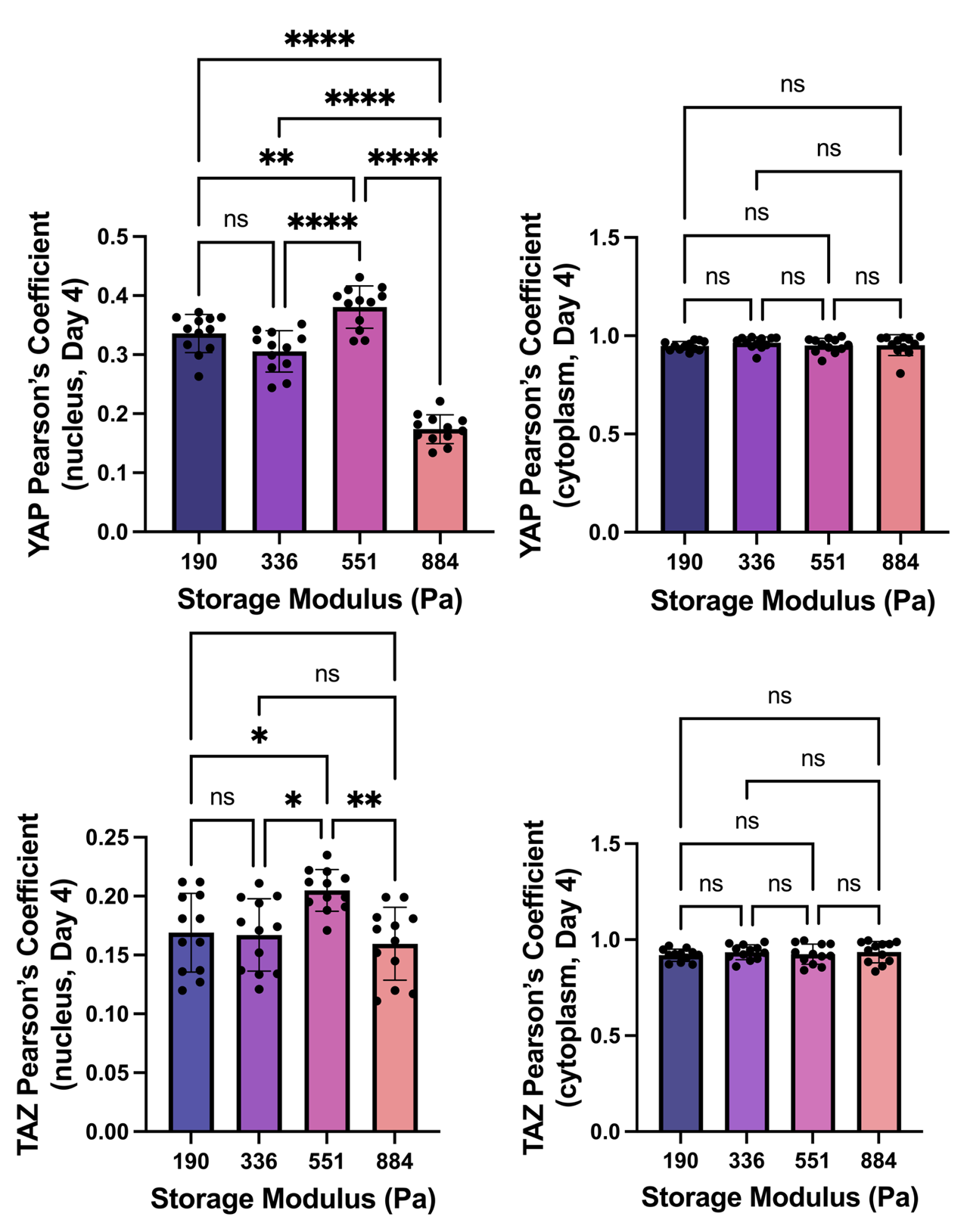

**Supplementary Figure 3:** Pearson’s correlation analysis of YAP and TAZ localization in hiPSC-EPs encapsulated in NorHA hydrogels of varying stiffness (190–884 Pa) after 4 days of culture. Nuclear YAP and TAZ localization are slightly enhanced at intermediate stiffness (551 Pa), while markedly reduced at higher stiffness (884 Pa). Cytoplasmic localization of both proteins remains relatively constant across all conditions. These results indicate that matrix stiffness modulates mechanotransduction signaling in encapsulated hiPSC-EPs, with nuclear translocation of YAP and TAZ serving as a marker of their activation. Statistical significance: ns (not significant), *p < 0.05, **p < 0.01, ****p < 0.0001 (one-way ANOVA with Tukey’s post hoc test).

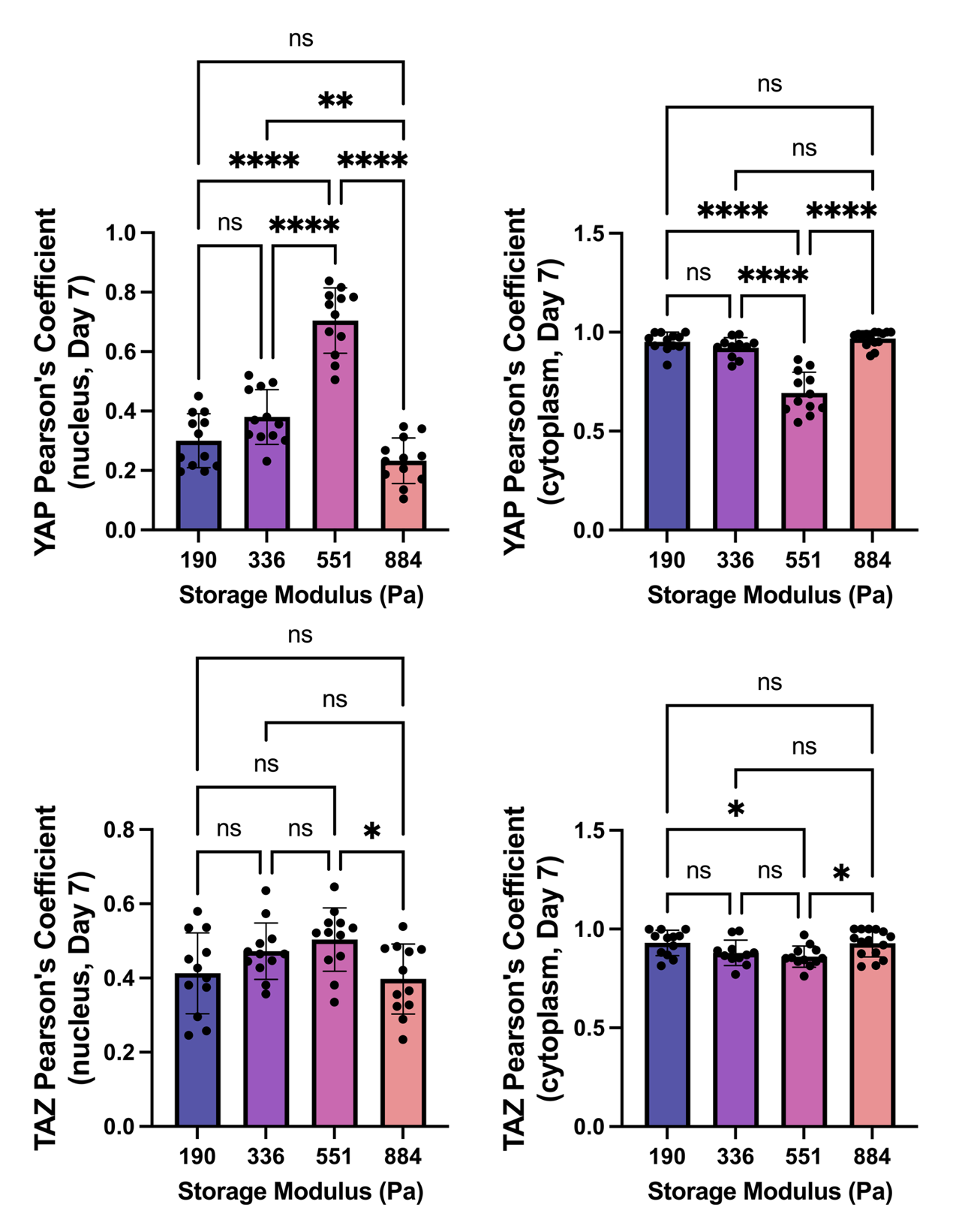

**Supplementary Figure 4:** **Extended culture enhances stiffness-dependent YAP nuclear localization in hiPSC-EPs encapsulated in NorHA hydrogels of varying stiffness (190–884 Pa). Pearson’s correlation analysis at day 7 reveals enhanced YAP nuclear localization in hiPSC-EPs encapsulated in 551 Pa hydrogels, showing a two-fold increase compared to 190 Pa hydrogels, with a significant reduction in 884 Pa hydrogels. Compared to day 4, nuclear YAP localization nearly doubled in the 551 Pa hydrogels. The enhanced nuclear-to-cytoplasmic YAP contrast at day 7 indicates progressive mechanotransduction signaling with extended culture time, maintaining the stiffness-dependent response pattern observed in day 4 cultures. Statistical significance: ns (not significant), *p < 0.05, **p < 0.01, ****p < 0.0001 (one-way ANOVA with Tukey post-test).**

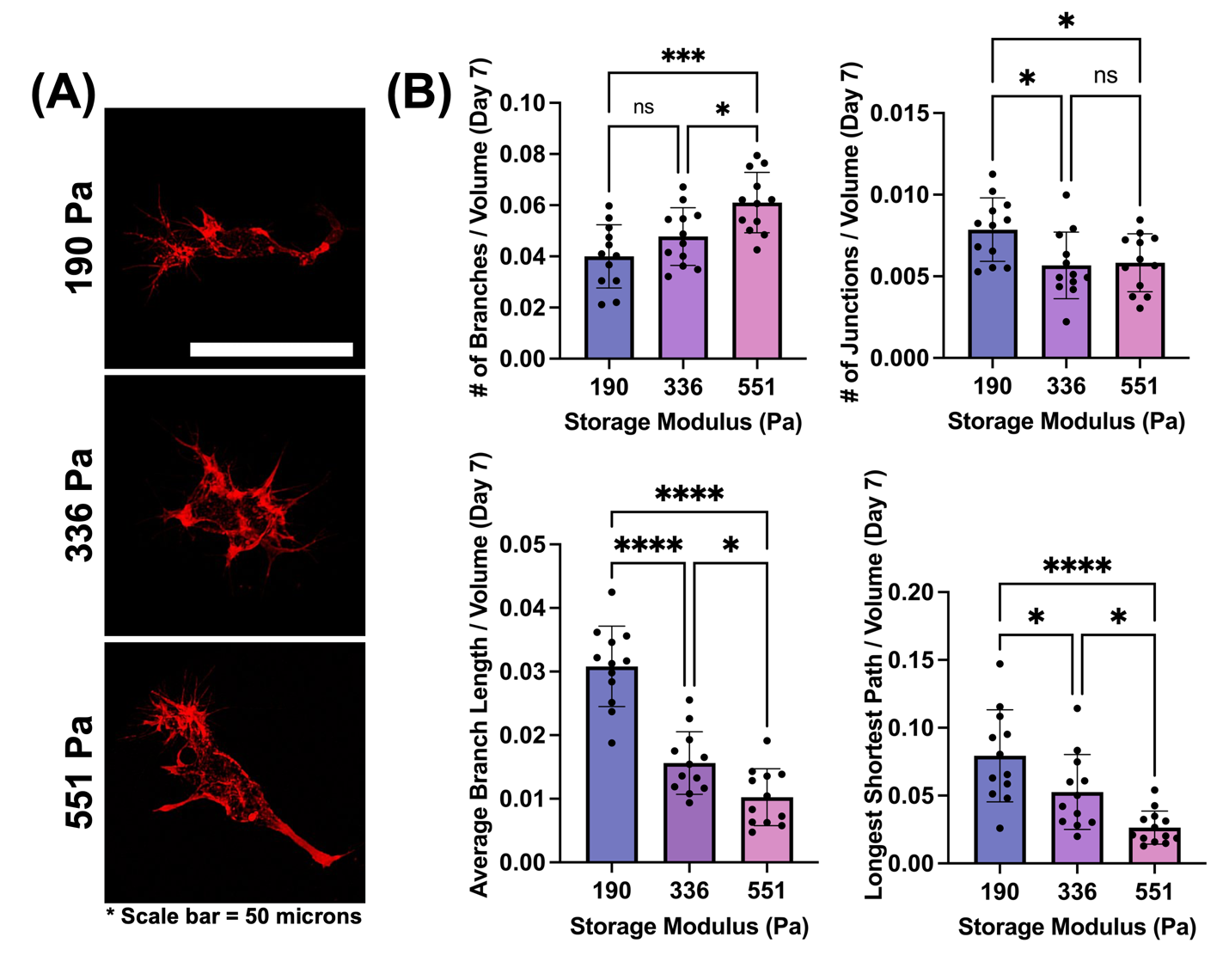

**Supplementary Figure 5:** Hydrogel stiffness modulates actin cytoskeleton architecture in hiPSC-EPs following 7 days of culture. (A) Confocal images of F-actin (phalloidin, red) in hiPSC-EPs encapsulated in hydrogels of varying stiffness (190, 336, and 551 Pa). hiPSC-EPs exhibit distinct morphological adaptations across the different hydrogel stiffnesses. Scale bar = 50 microns. (B) Quantification of F-actin network complexity. hiPSC-EPs cultured in the 551 Pa hydrogel exhibit the highest number of branches per volume, but the lowest branch length and shortest path metrics compared to complaint hydrogels. Notably, hiPSC-EPs cultured in the 190 Pa hydrogel show significantly higher average branch length per volume and longest shortest path per volume compared to both 336 Pa and 551 Pa conditions. Junction density appears slightly higher in the 190 Pa condition. Data presented as mean with individual data points; *P ≤ 0.05, ***P ≤ 0.001, ****P ≤ 0.0001, ns = not significant (one-way ANOVA and Tukey post-test).
