## Supplementary Table for "Matrix Stiffness Regulates Mechanotransduction and Vascular Network Formation of hiPSC-Derived Endothelial Progenitors Encapsulated in 3D Hydrogels"

| Solutions/Chemicals | | | Producer/Company | Product# | | Final Concentration |
| --- | --- | --- | --- | --- | --- | --- |
| Norbornene Hyaluronic Acid (NorHA, 52.5% functionalization, 445.536 g/mol) | | | Rosales Research Group (UT Austin) | N/A | | 10 mg/mL  (22.44 mM) |
| Endothelial Cell Growth Medium 2 (EGM-2) | | | Promo Cell | C-22011 | | N/A |
| LAP (lithium phenyl-2,4,6-trimethylbenzoylphosphinate) | | | Sigma-Aldrich | 900889-1G | | 0.025 wt% |
| DEG Peptide | Crosslinker Concentration  (% binding sites occupied) | Stiffness | Rosales Research Group (UT Austin) | N/A | - | |
|  | 25% | 190 Pa |  |  | 1.458 mM | |
|  | 50% | 336 Pa |  |  | 2.916 mM | |
|  | 75% | 551 Pa |  |  | 4.375 mM | |
|  | 100% | 884 Pa |  |  | 5.833 mM | |
| RGD Peptide | | | Rosales Research Group (UT Austin) | N/A | | 2 mM |
| Human VEGF 165 Protein | | | Acro Biosystems | VE5-H4210 | | 50 $n$g/mL |
| Y-27632 (ROCK) | | | Selleck Chemicals | S1049 | | 10 $\mu$M |

Supplementary Table 1: List of Materials Used for NorHA Hydrogel Formulation

| Solutions/Chemicals | Company | Product# | Final Concentration |
| --- | --- | --- | --- |
| DMEM/F12 | Sigma-Aldrich | D8437 | N/A |
| Essential 8 Medium | Thermo Scientific | A1517001 | N/A |
| Y-27632 (ROCK) | Selleck Chemicals | S1049 | 10 $\mu$M |
| B-27 Supplement, Insulin Free | ThermoFisher | A1895601 | 50x |
| N2 Supplement | ThermoFisher | 17502048 | 100x |
| Recombinant Human/Mouse/Rat Activin A | R&D Systems | 338-AC-050 | 25 $n$g/mL |
| Recombinant Human BMP4 Protein | R&D Systems | 314-BP-050 | 30 $n$g/mL |
| BIO | Selleck Chemicals | S7198 | 150 $n$M |
| SB 431542 | Selleck Chemicals | S1067 | 2 $\mu$M |
| Human VEGF 165 Protein | Acro Biosystems | VE5-H4210 | 50 $n$g/mL |

Supplementary Table 2: List of Materials Used for hiPSC-EP Differentiation
